# Neuropathological assessment of cerebrovascular integrity in Chronic Traumatic Encephalopathy

**DOI:** 10.64898/2026.11.09.765492

**Authors:** Adam D. Martinac, Anthony De Lorenzo, Derya Dik, Liana Pasas, Lynne E. Bilston, Claire E. Shepherd

**Affiliations:** Neuroscience Research Australia, Sydney 2031, Australia; Faculty of Medicine and Health, School of Biomedical Sciences, University of New South Wales, Sydney, NSW 2052, Australia; School of Biomedical Engineering, University of New South Wales, Sydney, NSW 2052, Australia

**Keywords:** Chronic traumatic encephalopathy (CTE), perivascular space (PVS), cerebrovascular pathology, pericytes, string vessels, AQP4

## Abstract

Chronic traumatic encephalopathy neuropathological change (CTE-NC) is defined by perivascular deposits of phosphorylated tau, with lesions preferentially located at sulcal depths. We investigated whether CTE-NC is associated with structural changes to perivascular spaces (PVSs) or microvascular degeneration. Post-mortem middle frontal gyrus tissue from 11 CTE-NC cases and 11 age-matched controls was examined using immunohistochemistry. All CTE-NC cases exhibited pathognomonic lesions in this region. We quantified PVS width along with vessel lumen and PVS area, string vessel burden, pericyte density, and aquaporin-4 (AQP4) polarisation to the astrocytic endfoot.

Total PVS width did not differ significantly between the groups in grey matter (P = 0.543) nor white matter (P = 0.468). Neither PVS area (sulcal depth, P = 0.097; gyral crest, P = 0.645) nor PVS-to-lumen area ratio (sulcal depth, P = 0.454; gyral crest, P = 0.307) differed significantly. Similarly, there were no significant differences in total microvascular length (p = 0.693), string vessel burden (p = 0.128), pericyte density (p = 0.589), or AQP4 endfoot enrichment (p = 0.538). We therefore found no evidence of large-scale PVS enlargement, lumen narrowing, capillary degeneration, pericyte loss, or altered AQP4 endfoot enrichment in CTE-NC at the locations and spatial scales investigated.

Taken together, these findings suggest that the pathognomonic perivascular tau lesion is not associated with a classical small vessel degenerative phenotype in CTE-NC. However, that does not exclude more subtle molecular, functional, or ultrastructural changes at the interface between vessels, the perivascular space, and surrounding brain tissue.

## Introduction

Perivascular spaces (PVSs) are fluid filled structures that surround arterioles and venules. These spaces are central to the hypothesised ‘glymphatic’ system, a lymphatic-like system which may be responsible for the exchange of cerebrospinal fluid and interstitial fluid, helping to maintain cerebral homeostasis (Iliff et al., 2012). Changes in PVS size and structure have been associated with neurodegenerative pathologies, including Alzheimer’s disease (AD) (Mestre et al., 2018; Wardlaw et al., 2020) and Parkinson’s disease (PD) (Dik et al., 2026). Alterations in vessel lumen diameter have also been associated with AD neuropathology (Iadecola et al., 2013). It has been proposed that these vascular changes may contribute to the pathogenesis of AD by impairing cerebral blood flow and/or nutrient/waste exchange. Given these findings in other neurodegenerative diseases, whether comparable structural vascular changes are present in chronic traumatic encephalopathy neuropathological change (CTE-NC) remains unclear.

CTE-NC is a neurodegenerative disease often found in individuals with a history of repetitive head impacts (Stein et al., 2014; Bieniek et al., 2021; McKee et al., 2023). Consensus criteria define the pathognomonic lesion of CTE-NC as phosphorylated tau aggregates in neurons, with or without glial tau, around small blood vessels at the depths of cortical sulci (Bieniek et al., 2021). This distribution within the sulcal depth is biologically important because sulcal depths are thought to be regions of high mechanical stress concentration during head impacts (Iverson et al., 2019; Rosen et al., 2023; Turk et al., 2022; Flavin et al., 2023). This suggests a possible mechanistic link between repetitive mechanical loading, vascular injury, and the molecular triggers of tau pathology, although this remains to be proven directly.

Several lines of evidence suggest that vascular dysfunction may contribute to CTE-NC pathogenesis. The leakage of serum proteins from damaged microvasculature may activate microglia, thereby inducing neuroinflammation, which is proposed to drive the pathogenesis of CTE-NC (Kirsch et al., 2023). In a mouse model of tauopathy, soluble pathogenic tau has been shown to enter brain microvascular endothelial cells and drive cellular senescence and brain microvascular dysfunction (Hussong et al., 2023). Human brain tissue studies have also reported increased severity and distribution of cerebral amyloid angiopathy (Standring et al., 2019), vascular injury markers (Kirsch et al., 2023) and cerebrovascular density and branching in CTE-NC (Rosen et al., 2023). This raises the possibility that CTE-NC involves vascular or perivascular alterations, but the effects of CTE-NC on PVS geometry, vessel lumen size, and related neurovascular unit components have not been comprehensively characterised in the human brain tissue.

Given their close association with the neurovascular unit, PVSs provide an interface between blood vessels, astrocytic endfeet, pericytes, and surrounding brain tissue. Subtle alterations in these components could occur even in the absence of overt PVS enlargement or vessel narrowing. Astrocytic endfeet are integral to the neurovascular unit and blood brain barrier, regulating water transport via polarised aquaporin-4 (AQP4) and providing a structural basis for perivascular exchange. Altered AQP4 expression has been reported after blast traumatic brain injury, with pronounced changes at sulcal depths and grey/white matter interfaces (Braun et al., 2024), while the regional and perivascular localisation of AQP4 in CTE-NC remains less clearly defined.

Importantly, our previous work in Parkinson’s disease has identified alterations in the microvasculature occurring alongside astrocytic and perivascular changes, suggesting that these components of the neurovascular unit may be affected in parallel in neurodegenerative disease (Dik et al., 2026). Pericytes within the neurovascular unit modulate capillary network properties and are plausible contributors to microvascular responses after injury (Winkler et al., 2011). String vessels, which are acellular capillary remnants, are widely interpreted as histological markers of endothelial injury and failed microvascular repair in neurodegeneration (Brown W.R., 2010; Yang et al., 2015; Erdener and Dalkara, 2019) and may provide insight into whether capillary dysfunction or degeneration are features of CTE-NC.

Despite the prominence of perivascular tau in the sulcal depth in CTE-NC, it remains unclear whether this pathology is accompanied by measurable changes in the cerebrovasculature. We therefore examined PVS morphology, capillary structure, string vessel burden, pericyte density, and AQP4 localisation in post-mortem middle frontal gyrus from CTE-NC cases and age-matched controls. We hypothesised that CTE-NC would be associated with enlarged PVSs, reduced vessel lumen size, AQP4 mislocalisation and evidence of microvascular degeneration.

## Materials and Methods

### Case selection and brain region analysed

The study included *ex vivo* human brain tissue from 22 individuals: 11 cases with chronic traumatic encephalopathy neuropathological change (CTE-NC) and 11 neurologically and neuropathologically normal controls (Table 1) (McCann et al., 2022). Tissue was obtained from the Sydney Brain Bank under project-specific ethics approval (UNSW Human Research Ethics Committee #211010).

**Table 1.** Demographic data and neuropathological scores for CTE-NC and control groups.

|  | **Controls** | **CTE-NC** |
|---|---|---|
| **N** | 11 | 11 |
| **Male/female** | 6/5 | 10/1 |
| **Age at death (Years)** | 83 ± 7 | 80 ± 7 |
| **Post-mortem delay (Hours)** | 25 ± 11 | 46 ± 35 |
| **Disease duration (Years)** | 0 ± 0 | 7 ± 5 |
| **Plaque severity (A score/3) (Shepherd et al. 2019)** | 1.3 ± 1.0 | 1.8 ± 1.1 |
| **NFT severity (B score/3) (Harding et al., 2000)** | 0.6 ± 0.5 | 2.7 ± 0.6* |
| **Braak LB stage (Braak et al., 2003)** | 0 ± 0 | 2.2 ± 2.6* |
| **CVD severity score (Esiri et al., 1997)** | 0.6 ± 0.5 | 1.2 ± 1.0 |

Between-group comparisons were assessed using t-tests, Mann-Whitney U tests or chi-squared tests, as appropriate.
Values are presented as mean ± SDev.
NFT = neurofibrillary tangle
LB = Lewy body
CVD = cerebrovascular disease
\**P* < 0.001 CTE-NC versus controls

All CTE-NC cases met current established neuropathological diagnostic criteria for CTE-NC (McKee et al., 2023). As all our CTE-NC cases were aged, co-pathologies were common, as previously described (Shepherd et al., 2024). Details of the co-pathologies are shown in **Supplementary Table 1**. Pathology scores for tau (B score), amyloid-plaques (A score), Lewy body stage (Braak Lewy body stage) and cerebrovascular disease (CVD score) are provided in Table 1.

We examined the middle frontal gyrus because it has been identified as a commonly and relatively early affected region in CTE-NC (McKee et al., 2015; Shepherd et al., 2024; Babcock et al., 2025; Affleck et al., 2025). All CTE-NC cases showed perivascular p-tau lesions in the middle frontal gyrus. Within the middle frontal gyrus, sampling was carried out in the grey and white matter and in paired sulcal depth and gyral crest regions. This comparison was selected because the sulcal depths are vulnerable to impact related biomechanical stress and are the site of pathognomonic CTE-NC lesions, whereas the gyral crest provides a nearby anatomical comparator, rather than a region assumed to be entirely free of pathology (Arena et al., 2020; Murray et al., 2022).

Although a total of 11 CTE-NC and 11 control cases were included, not all analyses could be quantified in every tissue section owing to section quality. Consequently, the sample size varied between analyses. The number of cases included for each variable is reported in **Supplementary Table 2**.

### Tissue processing and sectioning

Coronal formalin-fixed tissue blocks (3mm thick) were sampled from the middle frontal gyrus taking care to sample the sulcal depth and gyral crest. The blocks were embedded in paraffin and 12 serial 10µm thick sections were cut and dried onto Trajan Series 2 Adhesive Microscope Slides (Trajan Series 2 adhesive microslides 472042491 Scientific and Medical, Australia). Sections were deparaffinised using xylene and graded alcohols, then processed using microwave antigen retrieval (sections boiled for three minutes in 0.1M citrate buffer) for staining, as described below. **Supplementary Table 3** provides detailed information on all the antibodies used and their corresponding targets. For the negative controls, the primary antibody was omitted to confirm there was no non-specific binding of the secondary antibodies. No immunoreactivity was observed in these negative control sections.

### Peroxidase Immunohistochemistry

All the staining protocols used in this study have been previously published (Dik et al., 2026). Briefly, peroxidase immunohistochemistry using an antibody against collagen IV to detect the basement membrane was performed **(Supplementary Table 3)**. Prior to use, tissue sections were pre-treated with 1% H2O2 in 50% ethanol for 30 minutes and 10% normal horse serum for 20 minutes at room temperature. The primary antibody was incubated on the tissue sections overnight at 4°C. The sections were washed and incubated with their corresponding biotinylated secondary antibodies **(Supplementary Table 3)** for 30 minutes at 37°C. The sections were then washed and incubated with VECTASTAIN avidin-biotin tertiary antibody complex for 30 minutes at room temperature, visualised with 3,3’-diaminobenzidine (DAB) (Vector Laboratories, SK4105). The sections were then dehydrated through a series of graded alcohols and xylene, and cover-slipped using Dibutylphthalate Polystyrene Xylene (DPX). It is important to note that sections incubated with collagen IV were not counterstained, as a clean background was required to detect the capillary network and identify string vessels.

### Immunofluorescence

Perivascular spaces were identified by immunofluorescence using antibodies to detect vascular endothelial cells (rabbit anti-Factor VIII) and astrocytes (goat anti-GFAP). Smooth muscle actin (mouse anti-SMA-Cy3) was also used to differentiate between arterioles and venules to assess potential differences in PVS between these vessel types (see **Supplementary Table 3** for details). Tissue sections were incubated at 37°C for two hours, washed, and then incubated overnight at 37°C in the dark with secondary fluorophores corresponding to anti-Factor VIII and anti-GFAP. Smooth muscle actin (SMA) was conjugated to Cy3 and hence a secondary detection antibody was not required.

To examine the capillary network and pericytes, sections were stained with goat anti-collagen IV and counterstained with 4′,6-diamidino-2-phenylindole (DAPI), see Supplementary Table 2). Tissue was incubated overnight at 4°C with the primary antibody, washed, then incubated with the corresponding secondary fluorophore for 1 hour at room temperature in the dark.

To assess AQP4 localisation to astrocyte endfeet, sections were stained with rabbit anti-AQP4 and counterstained with DAPI (see **Supplementary Table 3** for details). Tissue sections were incubated overnight at 4°C with the primary antibody, washed, and then incubated with the corresponding secondary fluorophore for 1 hour at room temperature in the dark.

For all sections, autofluorescence was quenched by TrueBlack in 70% ethanol for five minutes, and the sections washed thoroughly prior to incubating with DAPI solution for at least one and a half hours in the dark at room temperature. Sections were then washed and coverslipped with an aqueous mounting medium.

### Image acquisition and region-of-interest selection

Images of whole tissue sections were acquired using an Olympus VS200 Research Slide Scanner. Brightfield and immunofluorescent sections were first scanned with a 2.5× objective to visualise overall anatomy and identify regions of interest (ROIs). High-resolution images for quantitative analysis were then acquired with a 20× objective. Within each tissue section, ROIs were sampled from predefined anatomical zones using either QuPath (Queen’s University Belfast, version 3.2) or Fiji/ImageJ (version 2.14.0/1.54f). Each ROI corresponded to a field of approximately 0.32 mm² (561 μm × 561 μm). Assay-specific ROI numbers and inclusion criteria are described in the relevant quantification sections below. The sampling scheme and metrics for each quantitative analysis are summarised in **Table 2**.

**Table 2.** Summary of quantitative histological analyses.

| **Analysis** | **Markers/stain** | **Regions sampled** | **Sampling** | **Metric** |
|---|---|---|---|---|
| **PVS width** | Factor VIII, SMA, GFAP | Grey matter and adja-cent white matter | 3 arterioles + 3 venules per region/case | Mean vessel-to-astrocytic boundary distance |
| **PVS area** | Factor VIII, GFAP | Sulcal depth, gyral crest | 5 vessels per region/case | PVS area, lumen area, PVS:lumen ratio |
| **String ves-sels** | Collagen IV, DAB | Grey matter | 5 ROIs/case | String vessel length / total capillary length |
| **Pericytes** | Collagen IV, DAPI | Grey matter | 5 ROIs/case | Pericytes per µm capillary length |
| **AQP4 local-isation** | AQP4, DAPI | Grey matter | 6 ROIs/case | Perivascular AQP4 / total AQP4 intensity |

PVS = perivascular space
AQP4 = aquaporin-4
SMA = smooth muscle actin
GFAP = glial fibrillary acidic protein
DAB = 3,3’-diaminobenzidine
ROI = region of interest

### Quantification of perivascular space

PVSs were identified in triple-labelled immunofluorescent sections using factor VIII to define the inner endothelial boundary of the vessel and GFAP to define the surrounding astrocytic boundary as previously described (Dik et al. 2026). SMA immunoreactivity was used to aid vessel classification, with arterioles identified by their strongly SMA positive vessel walls. PVS width was assessed by quantifying the mean distance between the vessel wall and astrocytic boundary in 3 arterioles and 3 venules in the grey matter and the subcortical white matter, yielding a total 18 vessels per case. Blood vessels sampled for analysis ranged from approximately 20–60 µm in cross-section. The images were exported into Fiji/ImageJ (version 2.14.0/1.54f), and the relevant objects were detected through pixel intensity on their corresponding fluorescent channels. For this analysis, the inner endothelial boundary and the outer astrocytic boundary of the vessel were the objects of interest. Manual deselection was used to remove non PVS objects. Two separate mask-like images were created for each vessel using the endothelial and astrocytic markers. A distance map was then generated in which each pixel was assigned a value based on the distance between the two boundaries, and the mean distance between the two objects was determined using a custom Fiji plug-in. Venule and arteriole mean vessel values were also summed per case to provide a combined mean distance value (total PVS, **see Figure 1**). The quantitative workflow is shown in **Supplementary Figure 1.**

**Figure 1.**
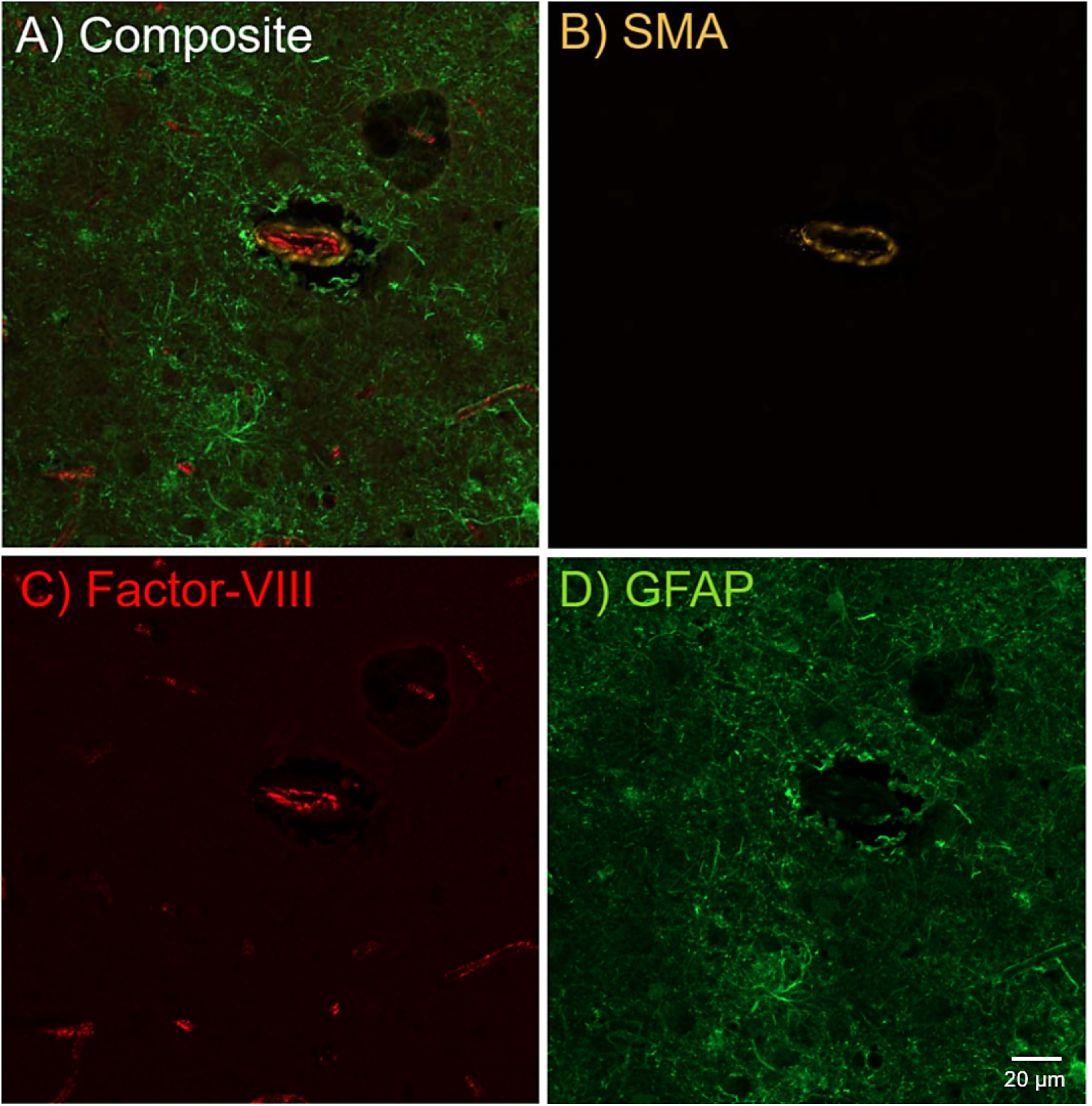
Representative image of a vessel with perivascular space showing a composite **(A**) of all fluorescent channels and each channel individually (B-D). (B/Yellow) SMA, (C/Red) Factor VIII, and (D/Green) GFAP. The scale bar is 20 microns.

For validation of this method, as well as the quantitative methods below, PVS structures in five randomly selected images from two different cases were independently counted by two researchers. Values varied by less than 5% between researchers, indicating high inter-rater reliability.

The PVS in the sulcal depth (SD) and the gyral crest (GC) was also assessed using a confirmatory area-based analysis. Vessel-centred ROIs were sampled from the SD and GC, with five vessels analysed per region per case. Only vessel profiles orientated approximately perpendicular to the imaging plane were included to minimise the effect of vessel orientation on cross-sectional area measurements. Blood vessels sampled for analysis ranged from approximately 15-30 µm in cross-section. Vessel type (arteriole or venule) was not stratified in this analysis, and vessel profiles were therefore analysed as a combined group.

ROIs selected in QuPath were imported into MATLAB for threshold assisted manual segmentation and area calculation. Because fluorescence intensity and background varied between staining batches, the GFAP and factor VIII channels were processed separately using background correction, mild smoothing and adaptive intensity thresholding to improve local contrast and aid identification of the relevant boundaries. Processing parameters were adjusted modestly where required to account for batch dependent differences in fluorescence intensity and background, rather than applying a single fixed threshold across all images. The processed images were then used to guide manual delineation of the outer GFAP defined perivascular boundary and the factor VIII defined vessel lumen, with the corresponding unprocessed fluorescence images retained as a reference during segmentation. Preprocessing therefore assisted identification and placement of the contours, but the final boundaries were defined manually rather than generated automatically from the thresholded pixels **(Supplementary Figure 2)**.

The two manually defined contours were converted to binary masks. The outer mask represented the total cross-sectional area enclosed by the GFAP-defined perivascular boundary, including the vessel lumen, while the inner mask represented the vessel lumen. PVS area was calculated by subtracting lumen area from the area enclosed by the outer boundary (**Supplementary Figure 2D**).

Three measurements were obtained for each vessel: lumen area, PVS area and the PVS:lumen area ratio, where 1 pixel = 0.119 µm². The PVS:lumen ratio was calculated as PVS area divided by lumen area, providing a measure of perivascular compartment size relative to vessel calibre rather than absolute PVS area alone. Vessel-level measurements were averaged within the SD and GC for each case before statistical analysis.

### Quantification of normal capillaries and capillary string vessels

Capillaries and string vessels were analysed in 5 randomly selected Regions of Interest (ROIs). Capillaries were defined as continuous collagen IV–positive tubular profiles forming a patent or skeletonisable lumen within the microvascular network (external diameter typically <10–12 µm). String vessels were defined as collagen IV–positive acellular basement-membrane remnants (non-lumenised filaments or short fragments) that either branch from, or lie between, adjacent capillaries and lack endothelial features.

Detection used QuPath for vessel masking and skeletonisation of the capillary network, with manual annotation of string vessels to avoid classifier false positives; obvious artefacts (tears, wrinkles, precipitate) were excluded. QuPath was unable to distinguish between normal capillaries and string vessels, making it necessary to manually segment normal capillaries from their branching string vessels using a QuPath annotation tool. Briefly, the ‘brush’ tool was used to isolate the string vessels from the normal capillaries using colour intensity (DAB peroxidase) (Dik et al., 2026). All annotations were then exported into Fiji using a custom QuPath Groovy script to analyse total length and to determine the proportion attributable to string vessels. Total capillary length and string vessel burden were averaged per case, and these values were used for statistical analysis. String vessel burden was expressed as cumulative string vessel length normalised to total capillary length within each ROI. The quantitative workflow is shown in **Supplementary Figure 3.**

### Quantification of pericytes

Pericytes were identified according to strict morphological criteria based on Ding et al. (2020). Specifically, the pericyte appeared as a protrusion lining the outer capillary wall and was surrounded by a collagen IV positive ring (membrane) with a clearly labelled DAPI-positive nucleus. Nucleated cell bodies located on the luminal side of the capillary wall were not included, as these were considered more likely to represent endothelial cells. Because pericytes were not observed on string vessel profiles, pericyte density was quantified only on normal capillaries, as previously described (Dik et al., 2026). The number of pericytes was counted in each sampled field and averaged per case for statistical analysis (pericytes per µm capillary length was used as the normalisation metric). The quantitative workflow is shown in **Supplementary Figure 4.**

### Quantification of aquaporin-4 polarisation to astrocytic endfeet

AQP4 polarisation to astrocytic endfeet was quantified in six ROIs per case. Blood vessels sampled for analysis ranged from approximately 20-60 µm in cross-section. Because arteriolar and venular PVS measures showed similar group effects (see Results), vessel type was not stratified for the AQP4 analysis. A quantitative approach adapted from Zeppenfeld et al. (2017) and Dik et al., (2026) was used to determine the amount of AQP4 polarised to perivascular astrocytic endfeet relative to total AQP4 signal within the sampled field. The fluorescent channel corresponding to AQP4 was selected, and the mean grey intensity of the entire ROI was measured as an index of total AQP4 expression. DAPI was used as a confirmatory stain to verify the identity of vascular profiles meeting the size criteria. A 10 µm perivascular border was drawn around each selected blood vessel, and the mean grey intensity of AQP4 immunofluorescence within this border was determined. The relative amount of AQP4 polarised to astrocytic endfeet was then calculated as the ratio of perivascular-border AQP4 intensity to total AQP4 intensity within the ROI. Vessel-level values were averaged per case for statistical analysis. The quantitative workflow is shown in **Supplementary Figure 5.**

## Statistical analysis

Statistical analyses were performed using IBM SPSS Statistics v.29 and GraphPad Prism v.10.2.3. A p-value of <0.05 was considered statistically significant. Group differences in demographics were analysed using t-tests and Mann Whitney U tests. Group differences in cerebrovascular disease (CVD), A and B score were assessed using chi-square tests. Group (CTE-NC versus controls) was included as a fixed factor in all subsequent models. Multiple regression analysis was undertaken to assess cellular and structural components associated with the capillary network, including string vessels and pericytes, with age and post-mortem delay included as covariates. Univariate regression models were performed to assess differences in PVS size between the CTE-NC and control group. Analyses were conducted for arteriolar and venular PVS size in the grey and white matter, total grey and white matter PVS size (representing the combined arteriolar and venular measures) and PVS area within the sulcal depth and gyral crest. Age, post-mortem delay and CVD were included as covariates for all models related to PVS. An additional univariate regression analysis was undertaken to assess group differences in AQP4 localisation to astrocytic endfeet. Age, postmortem delay and PVS size were included as covariates.

## Results

### Demographic and pathological differences between groups

The average age of the CTE-NC group was 80 ± 7 years, with an average post-mortem delay of 46 ± 35 hours. The average age of the control group was 83 ± 7 years, with an average post-mortem delay of 25 ± 11. (Table 1). There were no significant differences in age at death (Mann-Whiney U test, p = 0.300) or post-mortem delay (Mann-Whitney U test, p = 0.151) between the groups.

There was no significant difference in A score [χ^2^(3) *P* = 0.307] or CVD score [χ^2^(3) *P* = 0.175] between the CTE-NC and control group. B score was significantly different between the groups, reflecting increased tau pathology burden in CTE-NC [χ^2^(3) *P* < 0.001]. Braak Lewy body stage was significantly higher [χ^2^(1) *P* <0.017] in the CTE-NC group, consistent with co-pathology in this group (Supplementary Table 1). Demographic and neuropathological information of the cases used are given in **Table 1**.

### Perivascular space size is not enlarged in CTE-NC

There were no significant differences in PVS size between CTE-NC and control groups for either arterioles or venules in the grey matter (arterioles, p = 0.795; venules, p = 0.353) or white matter (arterioles, p = 0.252; venules, p = 0.805) (**Figure 2E–F; Supplementary Table S2**). Mean arteriolar PVS width was greater than mean venular PVS width in both groups. To assess whether the relative PVS widths of the two vessel types differed between CTE-NC and controls, we calculated the arteriolar-to-venular PVS width ratio. No significant between group differences were observed in either the grey matter (CTE-NC: 1.2 ± 0.28; controls: 1.3 ± 0.28; p = 0.191) or white matter (CTE-NC: 1.4 ± 0.28; controls: 1.7 ± 0.28; p = 0.086; Supplementary Table S2).

**Figure 2.**
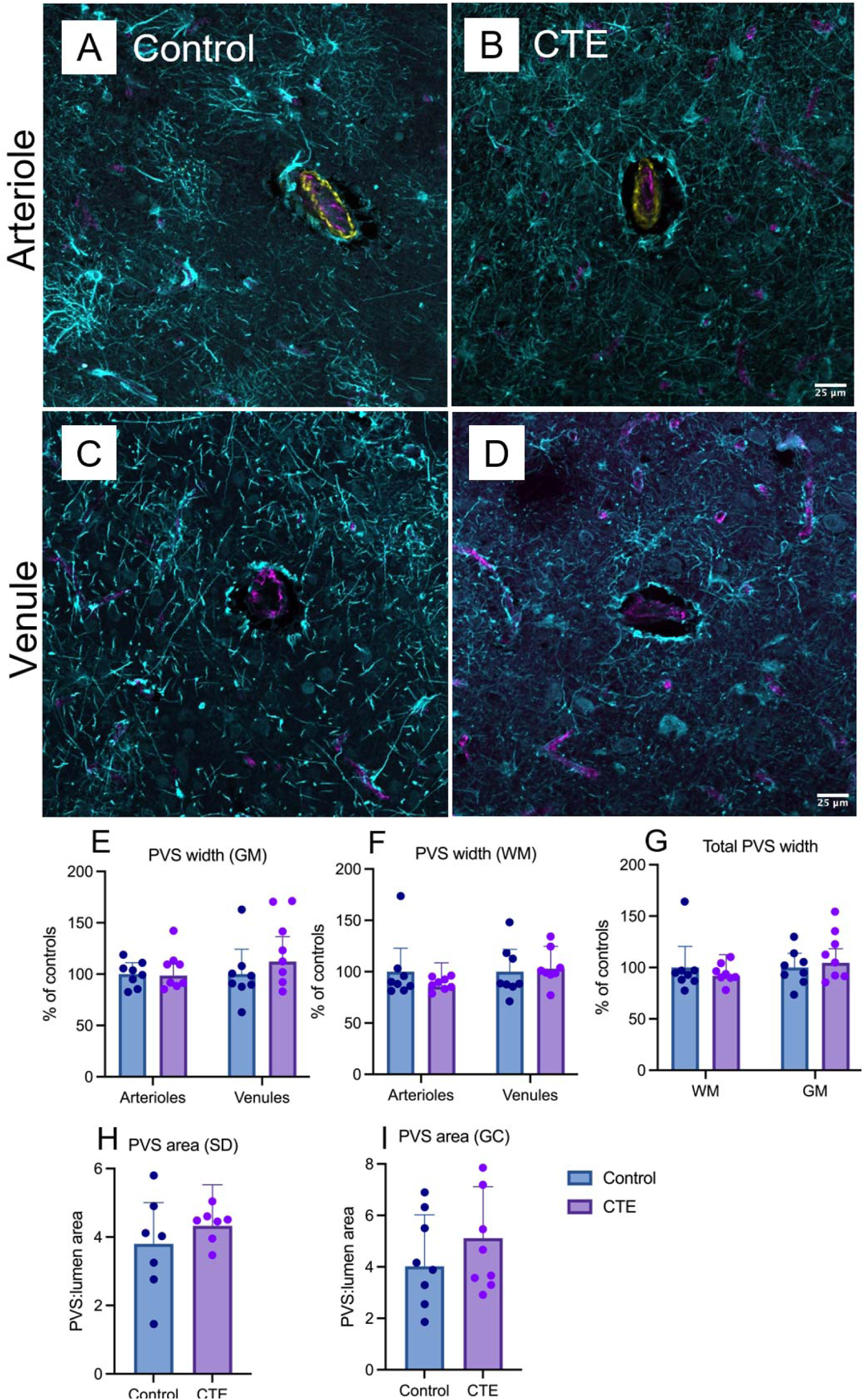
Perivascular space dimensions are preserved in CTE-NC. Representative immunofluorescence images of middle frontal gyrus grey matter from a control case **(A, C, left column)** and a CTE-NC case **(B, D, right column)** showing penetrating cortical arterioles (A, B) and venules (C, D) and associated perivascular space surrounded by astrocytic processes. Arteriole walls are labelled for smooth muscle (yellow), astrocytes are labelled with GFAP (cyan), and endothelial cells lining the vessel lumen (magenta). Bar graphs show **(E)** PVS width in arterioles and venules in the grey matter and **(F)** white matter, and total PVS width **(G)**, expressed as percentage of the controls. **(H)** PVS area at sulcal depth (SD) and **(I)** gyral crests (GC), expressed as the ratio of PVS to vessel lumen area. Blue bars indicate controls, and purple bars indicate CTE-NC cases and bars show group mean ± SDev. Individual points represent independent cases. PVS width did not differ significantly between control and CTE-NC in either the GM (p=0.543) or WM (p=0.468). The ratio between the PVS and lumen area did not vary between CTE-NC and control cases in the SD (p=0.454) or GC (p=0.307) Supplementary Table 2. The scale bar is 25 microns. CTE = chronic traumatic encephalopathy, PVS = perivascular space, GM = grey matter, WM = white matter, SD = sulcal depth, GC = gyral crest.

Since the vessel types did not differ between the CTE-NC and control group, arteriolar and venular measures were combined to determine whether there were overall differences between CTE-NC and control cases. There was no significant difference in the total PVS between the CTE-NC and control group in either the grey matter (p = 0.543) or the white matter (p = 0.468), see **Figure 2G**. Group means and statistical details are provided in **Supplementary Table 2.**

In terms of area metrics, there were no significant differences between CTE-NC and controls in PVS area, lumen area, or PVS:lumen area ratio at either the sulcal depth or gyral crest. At the sulcal depth, the mean PVS-to-lumen area ratio was 3.9 ± 1.2 in CTE-NC compared with 4.4 ± 0.51 in controls (Mann–Whitney U test, p = 0.383). The corresponding covariate-adjusted analysis also identified no significant between-group difference (p = 0.454; Supplementary Table S2). In the gyral crest, the mean PVS: lumen ratio was 4.3 ± 1.8 for CTE-NC and 4.8 ± 1.9 in controls (Figure 2H&I). Full regional means, standard deviations and p-values are reported in **Supplementary Figures 6-8**.

### Assessing the capillary network in CTE-NC

There were no significant differences in capillary network structure or capillary-associated pericyte density between CTE-NC and control cases. Capillary length did not differ significantly between groups (p = 0.693), and although string-vessel burden was numerically higher in CTE-NC cases than in controls (1.0% vs. 0.5%), this difference was not statistically significant (p = 0.128; **Supplementary Table 2**). Capillary associated pericyte density was not significantly different between CTE-NC and control groups (p = 0.589; Figure 3G–I). Group means and statistical summaries are provided in **Supplementary Table 2**.

**Figure 3.**
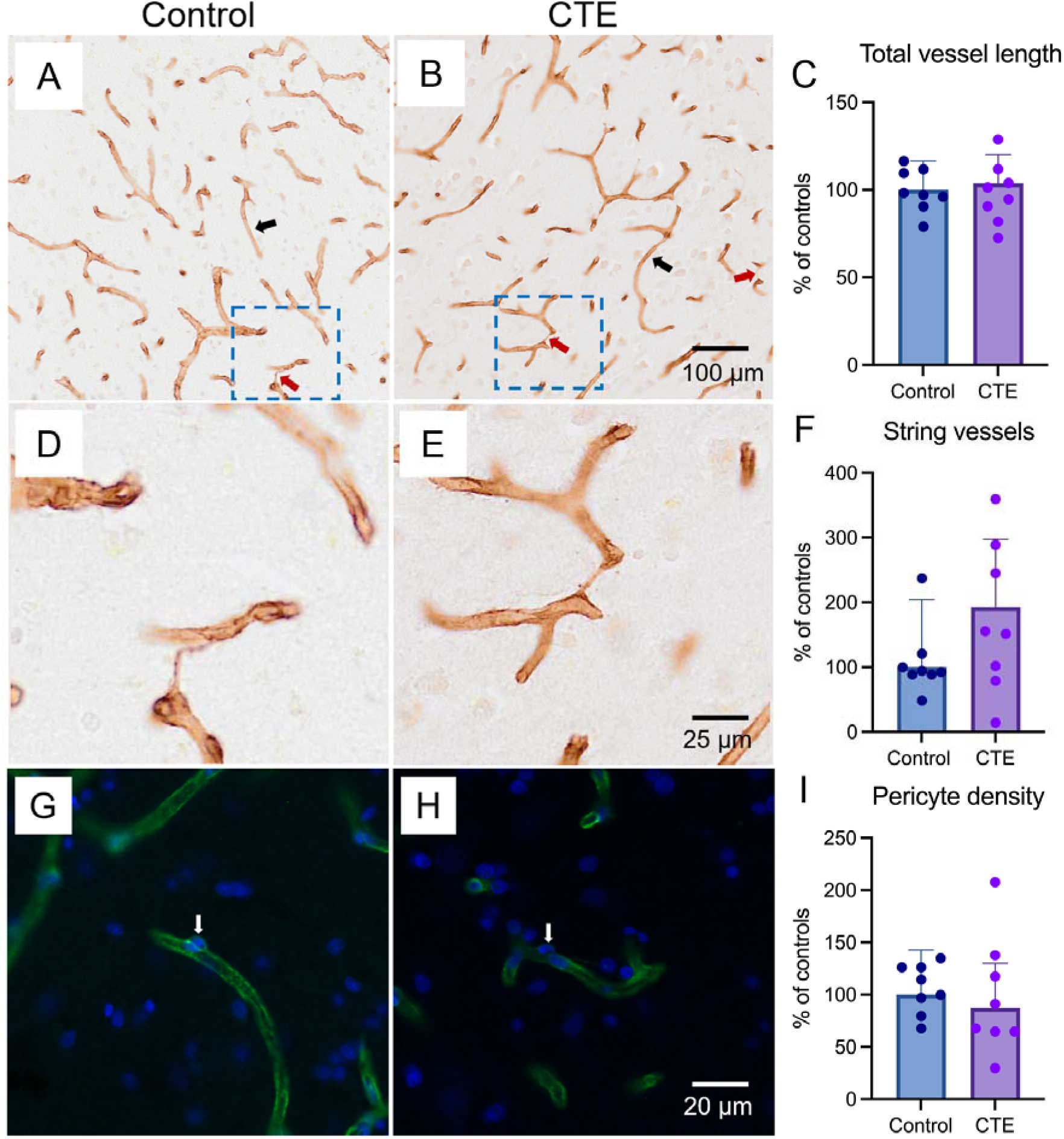
Total microvascular length, string vessel burden, and pericyte density in control and CTE-NC cortex. **(A, B)** Representative brightfield images of collagen IV–labelled microvessels in the middle frontal gyrus from a control case **(A)** and a CTE-NC case **(B).** Black arrows indicate intact capillaries; red arrows indicate string vessels. **(D, E)** Higher-magnification views from **(A, B)** illustrating thin, collapsed collagen IV–positive string vessels lacking an open lumen. **(C)** Quantification of total vessel length expressed as percentage of the control mean. **(F)** String vessel length normalised to total vessel length, expressed as percentage of the control mean. **(G, H)** Representative immunofluorescence images showing pericyte cell bodies (green; white arrow) along capillary walls and nuclei (blue) in control **(G)** and CTE-NC **(H)** cortex. **(I)** Pericyte density expressed as percentage of control pericytes. Bars and error bars show group mean ± SDev, individual points represent independent cases. Total vessel length (p = 0.693), string vessel burden (p = 0.128), and pericyte density (p = 0.589) did not differ significantly between control and CTE-NC groups. CTE-NC = chronic traumatic encephalopathy neuropathological change.

### Aquaporin-4 localisation to astrocytic endfeet in CTE-NC

There were no significant difference in AQP4 localisation between controls and CTE-NC (p = 0.538. **Figure 4C**). A summary of the group means, and significance can be found in **Supplementary Table 2**.

**Figure 4.**
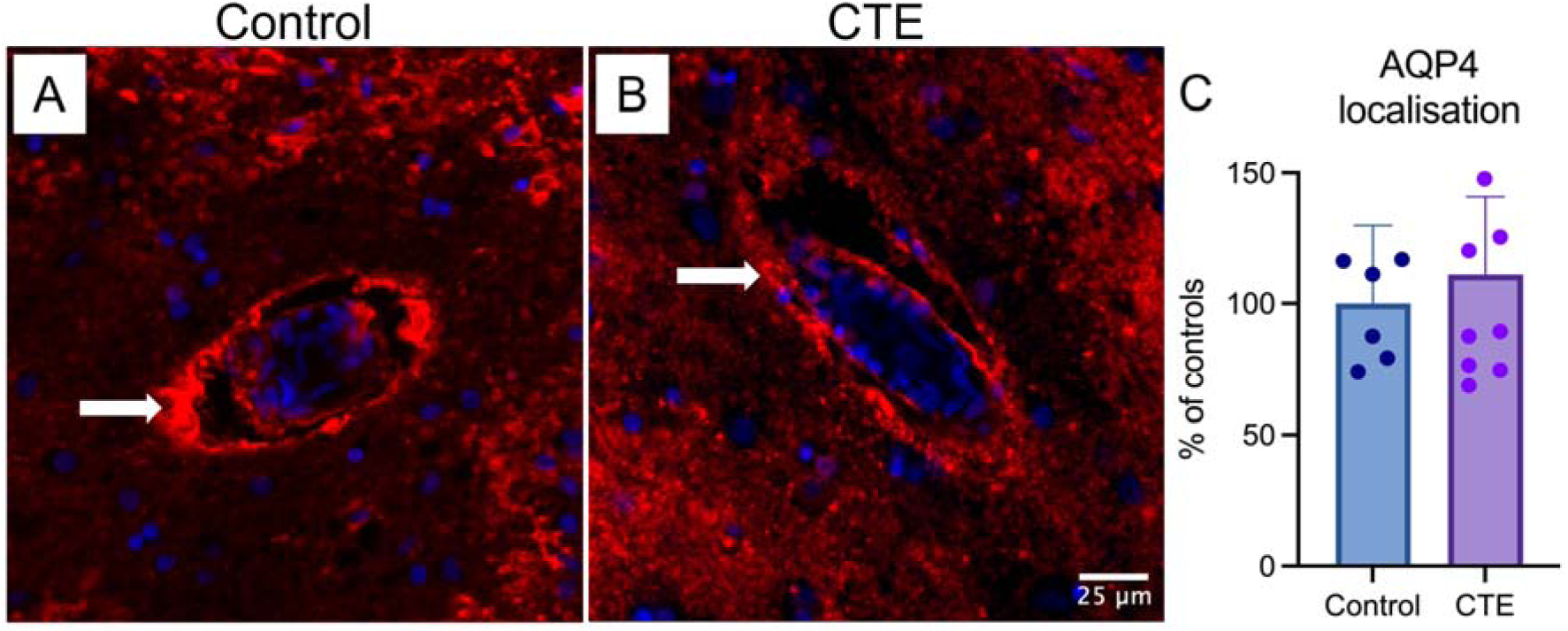
Perivascular aquaporin-4 polarisation is not significantly altered in CTE-NC compared with controls. **(A)** Representative immunofluorescence image of AQP4 (red) and DAPI-labelled nuclei (blue) around a blood vessel in control middle frontal gyrus tissue. The white arrow highlights AQP4 signal at the perivascular border. **(B)** Representative immunofluorescence image of AQP4 and DAPI around a blood vessel in CTE-NC middle frontal gyrus tissue, shown with the same measurement approach. Scale bar = 25 µm and applies to both **(A)** and **(B). (C)** Quantification of the perivascular endfeet AQP4 ratio, expressed as a percentage of the control mean. Bars and error bars show group mean ± SDev. Individual points represent independent cases. No significant difference was detected between CTE-NC and control groups (p = 0.538). CTE-NC = chronic traumatic encephalopathy neuropathological change, AQP4 = aquaporin-4.

## Discussion

This study quantified cellular changes in the PVS and microvasculature in the middle frontal gyrus in CTE-NC. Contrary to our initial hypotheses, no differences between CTE-NC cases and controls in PVS width, vessel lumen size, or the PVS:lumen area ratio were observed. The robustness of this finding was supported by two complementary morphometric approaches: the primary width-based analysis enabled separate assessment of arteriolar and venular PVSs, while the area-based analysis assessed PVS size relative to the associated vessel lumen. The absence of group differences was also consistent across grey and white matter and at the sulcal depth and gyral crest. Our findings in CTE-NC contrast with observations in AD and PD, where enlarged PVSs have been observed post-mortem and associated with increased disease severity and waste clearance dysfunction (Chen et al., 2022; Tu et al., 2022; Donahue et al., 2021; Ramirez et al., 2022; Lin et al., 2022; Dik et al., 2026). We also found no differences in string vessel burden, pericyte density, or AQP4 localisation between CTE-NC and control groups. Importantly, we have previously used the same analytical methods to demonstrate significant vascular alterations in PD (Dik et al., 2026), providing evidence that the methods are capable of detecting these changes when present. Overall, these findings indicate that overt structural alterations of the PVS and microvasculature are not a prominent feature of CTE-NC in the middle frontal gyrus, despite the presence of perivascular tau pathology.

### Perivascular spaces in CTE-NC

This is the first study to quantify the PVS architecture and associated cellular components in CTE-NC. Interestingly, we found no evidence of structural alteration of the arteriolar or venular PVS in CTE-NC cases compared to controls. In contrast, a recent in vivo MRI study of former American football players, who are a population at increased risk of CTE-NC (Daneshvar et al., 2026), reported an association between enlarged white matter PVS and greater lifetime exposure to repetitive head impacts (Jung et al., 2024). In addition, enlarged PVS volume was associated with worse cognitive and executive function, which the authors speculate may be due to impaired brain clearance (Jung et al 2024). However, other imaging studies have reported less pronounced or absent differences. For example, an MRI study comparing football players (high-contact) with volleyball players (low-contact) found no significant group differences in PVS (McAllister et al 2021). Similarly, Yamamoto and colleagues (2024) found no difference in PVS burden (volume and number) between former veterans with and without blast related brain injury, although PVS burden was associated with cerebrospinal fluid inflammatory markers within the head injury group, suggesting an association between central inflammation, head injury and the PVS (Yamamoto et al 2024). Furthermore, PVS volume has been associated with age in other populations (Barisano et al., 2021; Lynch et al., 2023; Kim et al., 2023), highlighting the potential influence of demographic and other factors on PVS measurements. Collectively, these findings suggest that the relationship between head-impact exposure and PVS abnormalities is complex and may vary according to population characteristics and the clinical context.

Evidence from broader traumatic brain injury populations provides additional context but should be interpreted separately from studies of sports related repetitive head impact, especially since isolated traumatic brain injury is not reliably associated with CTE-NC (McCann et al 2024). In a population of individuals with chronic traumatic brain injury, Hicks and colleagues (2023) reported a greater burden of enlarged PVS, particularly in individuals with bilateral brain lesions, with greater burden associated with poorer verbal memory performance. In mild traumatic brain injury, Zhuo et al. (2024) found that acute white matter enlarged PVS burden was associated with post-concussion symptoms, while longitudinal changes in the analysis along the perivascular space (ALPS) index were associated with sleep quality outcomes. The ALPS index should be interpreted cautiously as an indirect diffusion measure, rather than a direct measure of waste clearance function (Taoka et al. 2024). These studies indicate that PVS-related imaging measures may vary following traumatic brain injury, but their relevance to the neuropathological changes associated with repetitive head injury and CTE-NC remains uncertain.

MRI visible PVS and histologically measured PVS also differ in scale and measurement and may capture different elements of perivascular pathology. In particular, our assessment of small vessels (20-50µm) does not establish whether larger penetrating vessels and their associated PVS, which are more reliably detected by MRI, are similarly preserved. Our observation that PVS width and PVS:lumen ratios are preserved in the grey and white matter and at cortical sulcal depths and gyral crests in CTE-NC suggests that PVS-related imaging abnormalities do not necessarily correspond to persistent gross histological enlargement in CTE-NC. Thus, PVS-related imaging abnormalities reported in repetitive head injury and traumatic brain injury may not directly imply gross histological enlargement in CTE-NC. Whether these imaging findings reflect differences in vessel calibre, regional or temporal variation, or functional and molecular alterations without persistent structural changes remains to be seen.

### Vascular pathology beyond the PVS

String vessels, acellular remnants of regressed capillaries, are considered markers of severe endothelial injury, failed vascular repair, and capillary loss. They have been widely reported in neurodegenerative conditions such as AD (Challa et al., 2004; Brown, 2010; Steinman et al., 2021) and PD (Yang et al., 2015; Lau et al., 2024; Dik et al., 2026). In contrast, we found no meaningful increase in string-vessel burden in CTE. We also found no evidence of reduced capillary-associated pericyte density. Pericytes are key regulators of blood–brain barrier integrity and capillary blood flow (Armulik et al., 2010; Daneman et al., 2010; Hall et al., 2014), and their loss has been linked to endothelial degeneration and string-vessel formation in other neurodegenerative settings (Bell et al., 2010; Sengillo et al., 2013; Yang et al., 2015; Dik et al., 2026). The absence of increased string-vessel burden, together with preserved capillary-associated pericyte density, suggests that overt capillary regression is not a prominent feature of CTE-NC in the region examined.

This pattern appears distinct from the regressive microangiopathy described in AD and PD, where string-vessel accumulation and capillary rarefaction are more commonly reported (Erdener and Dalkara, 2019; Dik et al., 2026). Rosen et al. (2023), previously reported increased capillary branching in CTE-NC using 3D vascular reconstructions, particularly in the sulcal depth where associations with tau pathology were evident. The authors propose a role for angiogenesis in CTE-NC due to secondary hypoxia arising from impact-related vessel damage. They also highlight the potential influence of cortical atrophy and regional variation, which were not assessed in the present study, but may have affected the observed outcomes.

Two imaging studies have found reductions in cerebral blood in individuals exposed to repetitive head impacts, which may be related to capillary structure and pericyte number (Dik et al., 2026). Using arterial spin labelling MRI, Querzola reported reductions in cerebral blood flow in sensorimotor areas in American football players compared to healthy controls (Querzola et al., 2022) while Amen identified similarly reported lower perfusion in 36 out of 128 atlas-defined brain regions in current and former NFL players, with perfusion measures distinguishing players from healthy controls (Amen et al 2016). Importantly, these studies were conducted in living athletes exposed to repetitive head impacts, which is associated with an increased risk of CTE-NC, but CTE-NC was not neuropathologically confirmed. However, our findings indicate that pericyte density and string vessel burden were unchanged in the middle frontal gyrus, suggesting that altered cerebral perfusion, if present in CTE-NC, may reflect functional cerebrovascular changes that are not necessarily accompanied by overt microvascular degeneration. Altered neuronal activity may also explain changes in cerebral perfusion through altered neurovascular coupling, although this remains speculative.

Endothelial cell dysfunction, reduced cerebral blood flow, glymphatic impairment and tau accumulation have also been reported in mouse models of repetitive head injury (Iliff et al., 2014; Lynch et al., 2016; Ojo et al., 2016; Eisenbaum et al., 2021; Eisenbaum et al., 2024; Cembran et al., 2025). Furthermore, significant vascular inflammation, endothelial cell dysfunction and senescence have been identified in CTE-NC brain tissue, especially in cases with high levels of tau pathology (Eisenbaum et al. 2026). The authors suggest that endothelial cell degeneration is a prominent feature of gliovascular decline following repetitive head injury, leading to oxidative stress and hypoxia.

We found no difference in AQP4 localisation in the CTE-NC cases compared to controls (Fig 4). This finding doesn’t preclude alterations in total AQP4 abundance or more spatially restricted changes in AQP4 expression. However, we previously identified reduced perivascular AQP4 polarisation in PD despite relatively preserved total AQP4 expression (Dik et al., 2026), demonstrating the ability of this methodology to detect alterations in AQP4 localisation under other neuropathological conditions. Importantly, AQP4 changes have been reported within CTE-NC lesions. Osterman et al. (2025) observed increased AQP4 immunoreactivity around p-tau lesion vessels at the sulcal depth compared with adjacent non-lesioned tissue in the gyral wall. Although our study similarly compared sulcal and gyral regions, vessels were not selected based on their association with p-tau lesions. Given the focal distribution of CTE-NC pathology, this difference in sampling may have limited our sensitivity to lesion-specific alterations in AQP4. Furthermore, both Osterman et al. (2025) and Eisenbaum et al. (2026) used spatial approaches to investigate AQP4 distribution, whereas we quantified relative perivascular-to-global AQP4 enrichment. These methodological differences should be considered when comparing findings.

Recent studies have investigated changes in cerebrovascular endothelial activation and inflammatory signalling in CTE-NC. The intercellular adhesion molecule 1 (ICAM1) and vascular cell adhesion molecule 1 (VCAM1) are endothelial adhesion proteins that are upregulated during endothelial activation and inflammation, while C-reactive protein (CRP) is a marker of inflammatory response (Kirsch et al., 2023). Increased levels of ICAM1, VCAM1, and CRP have been reported in CTE-NC brain tissue compared with controls (Kaur et al., 2022; Kirsch et al., 2023). Increased ICAM1 immunoreactivity has also been described in the microvasculature, extracellular spaces, and astrocytes of CTE-NC affected brains, consistent with vascular injury and local inflammatory signalling (Kirsch et al., 2023). Importantly, these changes were observed in the dorsolateral prefrontal cortex, which is a functional region that includes the middle frontal gyrus, and occurred despite preserved vascular integrity, which is consistent with our current observations. Complementing these findings, spatial transcriptomic analysis has identified a common 21-gene signature associated with tau-positive lesions implicating astrocytic activation, neuroinflammation, blood brain barrier function, and extracellular matrix remodelling (Suter et al. (2025). Together, these findings suggest that CTE-NC may be associated with inflammatory and molecular changes in the cerebrovascular environment, which occur in the absence of overt structural alterations in the cerebrovasculature.

## Limitations

The area-based analysis relied on threshold-assisted manual segmentation, with modest adjustment of preprocessing parameters required to accommodate variation in fluorescence between staining batches. This introduces some operator dependence and could have reduced sensitivity to subtle differences in PVS area-based measures. However, no evidence of PVS enlargement was observed using methods that can detect PVS enlargement in PD (Dik et al 2026).

We assessed PVS width and area measures in fixed human post-mortem tissue, as is common in histological studies of perivascular structure. However, post-mortem fixation and tissue processing may alter PVS dimensions, and the extent to which PVSs collapse or shrink after death remains uncertain. Plá et al. (2023) showed that hyperosmotic fixatives can cause dehydration and shrinkage of meningeal membranes and associated perivascular spaces in mice, suggesting that some post-mortem tissue changes may be artefacts. Absolute PVS dimensions in fixed tissue may therefore not fully reflect *in vivo* geometry, and differences in tissue state may contribute to discrepancies between in vivo MRI studies reporting changes in PVSs after repetitive head impacts and the preserved PVS dimensions observed here. Because CTE-NC and control tissues were processed using comparable workflows, fixation differences are unlikely to explain the absence of group differences, especially since our technique is also able to detect PVS enlargement in PD (Dik et al 2026). However, our findings only address gross PVS enlargement in fixed post-mortem tissue and cannot exclude subtle or dynamic changes that occur *in vivo*.

We also acknowledge the small sample size of both our CTE-NC and control groups. Cases and controls were selected from the same brain bank to reduce variability related to tissue handling, fixation, and processing, which limits sample size. Another limitation is that pericytes were identified based on strict morphological criteria, rather than using a specific pericyte marker. Our approach is aligned with previous studies of pericyte loss in PD (Dik et al., 2026), which showed consistent results with earlier research (Yang et al., 2015). Although this approach allowed conservative identification of capillary-associated pericytes, the absence of a specific pericyte marker means that we could not assess biochemical or phenotypic changes in pericytes. Our results therefore address pericyte density rather than pericyte activation, dysfunction, or molecular state.

Another important consideration is the presence of coexisting neuropathologies in many of the CTE-NC cases, in particular many cases had concomitant Alzheimer’s and Lewy body pathology. This is relevant because cerebrovascular abnormalities have been reported in both AD and PD (Dik et al., 2026), raising the possibility that co-pathology could influence the vascular measures examined here. Importantly, despite this co-pathology, the CTE-NC cases in the present study did not show significant PVS enlargement, capillary regression, pericyte loss, or altered relative AQP4 localisation in the middle frontal gyrus.

## Conclusion

CTE-NC was not associated with overt PVS enlargement, vessel narrowing, capillary regression, pericyte loss, or altered relative AQP4 localisation in the middle frontal gyrus. These findings indicate that gross regional structural alterations of the PVS and microvasculature are not a prominent feature of CTE-NC, despite the presence of perivascular tau pathology. Whether localised molecular alterations or larger-calibre vessels and their associated PVS are structurally altered in CTE-NC remains an important question for future neuropathological studies.

## Declarations

C.E.S serves as a member of the steering committee for the Australian Football League Brain Health Initiative

## Acknowledgements

Tissues were received from the Sydney Brain Bank which is supported by Neuroscience Research Australia and a special gift in memory of Jim Raftos from the Shaw family. Adam Martinac was supported by the National Health and Medical Research Council (NHMRC) Ideas grant [APP2011940].

## References

1. Affleck, A. J., Suter, C. M., Cropley, J. E., Pearce, A. J., & Buckland, M. E. (2025). The neuropathology of chronic traumatic encephalopathy. Pathology, 57(2), 248–252. 10.1016/j.pathol.2024.12.387

2. Amen, D. G., Willeumier, K., Omalu, B., Newberg, A., Raghavendra, C., & Raji, C. A. (2016). Perfusion neuroimaging abnormalities alone distinguish National Football League players from a healthy population. Journal of Alzheimer’s Disease, 53(1), 237–241. 10.3233/JAD-160207

3. Arciniega, H., Wickham, A., Szekely, B., Kim, N., Cho, K. I., Carrington, H., Knyazhanskaya, E. E., John, O., Jung, L. B., Breedlove, K., Mirmajlesi, A. S., Stearns, J., Rushmore, R. J., Daneshvar, D. H., Wiegand, T. L. T., Billah, T., Pasternak, O., Cetin-Karayumak, S., Rathi, Y., … Bouix, S. (2026). Dose-dependent white matter changes associated with repetitive head impacts in former American football players. Brain Communications, 8(3), Article fcag195. 10.1093/braincomms/fcag195

4. Arena, J. D., Johnson, V. E., Lee, E. B., Gibbons, G. S., Smith, D. H., Trojanowski, J. Q., & Stewart, W. (2020). Astroglial tau pathology alone preferentially concentrates at sulcal depths in chronic traumatic encephalopathy neuropathologic change. Brain Communications, 2(2), Article fcaa210. 10.1093/braincomms/fcaa210

5. Armulik, A., Genové, G., Mäe, M., Nisancioglu, M. H., Wallgard, E., Niaudet, C., He, L., Norlin, J., Lindblom, P., Strittmatter, K., Johansson, B. R., & Betsholtz, C. (2010). Pericytes regulate the blood–brain barrier. Nature, 468(7323), 557–561. 10.1038/nature09522

6. Babcock, K. J., Abdolmohammadi, B., & McKee, A. C. (2025). Recent advances in chronic traumatic encephalopathy. The American Journal of Pathology, 195(11), 2048–2058. 10.1016/j.ajpath.2025.07.008

7. Barisano, G., Sheikh-Bahaei, N., Law, M., Toga, A. W., & Sepehrband, F. (2021). Body mass index, time of day and genetics affect perivascular spaces in the white matter. Journal of cerebral blood flow and metabolism : official journal of the International Society of Cerebral Blood Flow and Metabolism, 41(7), 1563–1578. 10.1177/0271678X20972856

8. Bell, R. D., Winkler, E. A., Sagare, A. P., Singh, I., LaRue, B., Deane, R., & Zlokovic, B. V. (2010). Pericytes control key neurovascular functions and neuronal phenotype in the adult brain and during brain aging. Neuron, 68(3), 409–427. 10.1016/j.neuron.2010.09.043

9. Bieniek, K. F., Cairns, N. J., Crary, J. F., Dickson, D. W., Folkerth, R. D., Keene, C. D., Litvan, I., Perl, D. P., Stein, T. D., Vonsattel, J. P., Stewart, W., Dams-O’Connor, K., Gordon, W. A., Tripodis, Y., Alvarez, V. E., Mez, J., Alosco, M. L., & McKee, A. C. (2021). The second NINDS/NIBIB consensus meeting to define neuropathological criteria for the diagnosis of chronic traumatic encephalopathy. Journal of Neuropathology & Experimental Neurology, 80(3), 210–219. 10.1093/jnen/nlab001

10. Braak, H., Del Tredici, K., Rüb, U., de Vos, R. A., Jansen Steur, E. N., & Braak, E. (2003). Staging of brain pathology related to sporadic Parkinson’s disease. Neurobiology of aging, 24(2), 197–211. 10.1016/s0197-4580(02)00065-9

11. Braun, M., Sevao, M., Keil, S. A., Gino, E., Wang, M. X., Lee, J., Haveliwala, M. A., Klein, E., Agarwal, S., Pedersen, T., Rhodes, C. H., Jansson, D., Cook, D., Peskind, E., Perl, D. P., Piantino, J., Schindler, A. G., & Iliff, J. J. (2024). Macroscopic changes in aquaporin-4 under-lie blast traumatic brain injury-related impairment in glymphatic function. Brain, 147(6), 2214–2229. 10.1093/brain/awae065

12. Brown, R., Benveniste, H., Black, S. E., Charpak, S., Dichgans, M., Joutel, A., Nedergaard, M., Smith, K. J., Zlokovic, B. V., & Wardlaw, J. M. (2018). Understanding the role of the perivascular space in cerebral small vessel disease. Cardiovascular Research, 114(11), 1462–1473. 10.1093/cvr/cvy113

13. Brown, W. R. (2010). A review of string vessels or collapsed, empty basement membrane tubes. Journal of Alzheimer’s Disease, 21(3), 725–739. 10.3233/JAD-2010-100219

14. Challa, V. R., Thore, C. R., Moody, D. M., Anstrom, J. A., & Brown, W. R. (2004). Increase of white matter string vessels in Alzheimer’s disease. Journal of Alzheimer’s Disease, 6(4), 379–383. 10.3233/JAD-2004-6404

15. Cembran, A., Eisenbaum, M., Paris, D., Mullan, M., Crawford, F., Ferguson, S., & Bachmeier, C. (2025). Contribution of brain pericytes to neuroinflammation following repetitive head trauma. Journal of Inflammation, 22, Article 11. 10.1186/s12950-025-00439-9

16. Chen, H., Wan, H., Zhang, M., Wardlaw, J. M., Feng, T., & Wang, Y. (2022). Perivascular space in Parkinson’s disease: Association with CSF amyloid/tau and cognitive decline. Parkinsonism & Related Disorders, 95, 70–76. 10.1016/j.parkreldis.2022.01.002

17. Daneman, R., Zhou, L., Kebede, A. A., & Barres, B. A. (2010). Pericytes are required for blood–brain barrier integrity during embryogenesis. Nature, 468(7323), 562–566. 10.1038/nature09513

18. Daneshvar, D. H., Nowinski, C. J., Abdolmohammadi, B., Luster, C. B., Uretsky, M., Martin, B. M., Palmisano, J. N., Weuve, J., McClean, M. D., Cherry, J. D., Dwyer, B., Feigel, E. D., Mastrodicasa, M. J., Alvarez, V. E., Rabinovici, G. D., Seeley, W. W., Grinberg, L. T., Crary, J. F., Stein, T. D., … Mez, J. (2026). Prevalence of chronic traumatic encephalopathy at death in National Football League players: Retrospective population based cohort study, 2008–21. BMJ, 394, Article e100418. 10.1136/bmj-2026-100418

19. Dik, D., Halliday, G. M., Sytnyk, V., & Shepherd, C. E. (2026). Region-specific variations in the cerebrovasculature underlie disease progression in Parkinson’s disease. Brain, 149(2), 592–605. 10.1093/brain/awaf305

20. Ding, R., Hase, Y., Ameen-Ali, K. E., Ndung’u, M., Stevenson, W., Barsby, J., Gourlay, R., Akinyemi, T., Akinyemi, R., Uemura, M. T., Polvikoski, T., Mukaetova-Ladinska, E., Ihara, M., & Kalaria, R. N. (2020). Loss of capillary pericytes and the blood–brain barrier in white matter in poststroke and vascular dementias and Alzheimer’s disease. Brain Pathology, 30(6), 1087–1101. 10.1111/bpa.12888

21. Donahue, E. K., Murdos, A., Jakowec, M. W., Sheikh-Bahaei, N., Toga, A. W., Petzinger, G. M., & Sepehrband, F. (2021). Global and regional changes in perivascular space in idiopathic and familial Parkinson’s disease. Movement Disorders, 36(5), 1126–1136. 10.1002/mds.28473

22. Eisenbaum, M., Pearson, A., Gratkowski, A., Mouzon, B., Mullan, M., Crawford, F., Ojo, J., & Bachmeier, C. (2021). Influence of traumatic brain injury on extracellular tau elimination at the blood–brain barrier. Fluids and Barriers of the CNS, 18(1), Article 48. 10.1186/s12987-021-00283-y

23. Eisenbaum, M., Pearson, A., Ortiz, C., Koprivica, M., Cembran, A., Mullan, M., Crawford, F., Ojo, J., & Bachmeier, C. (2024). Repetitive head trauma and apoE4 induce chronic cerebrovascular alterations that impair tau elimination from the brain. Experimental Neurology, 374, Article 114702. 10.1016/j.expneurol.2024.114702

24. Eisenbaum, M., Pearson, A., Cembran, A., Mufson, E. J., Ortiz, C., Spelman, M., Mullan, M., Crawford, F., Ojo, J., & Bachmeier, C. (2026). Mild traumatic brain injury promotes chronic cerebrovascular inflammation and glymphatic suppression. Molecular neurodegeneration advances, 2(1), 45. 10.1186/s44477-026-00053-w

25. Erdener, Ş. E., & Dalkara, T. (2019). Small vessels are a big problem in neurodegeneration and neuroprotection. Frontiers in Neurology, 10, Article 889. 10.3389/fneur.2019.00889

26. Esiri, M. M., Wilcock, G. K., & Morris, J. H. (1997). Neuropathological assessment of the lesions of significance in vascular dementia. *Journal of Neurology*, Neurosurgery & Psychiatry, 63(6), 749–753. 10.1136/jnnp.63.6.749

27. Flavin, W. P., Hosseini, H., Ruberti, J. W., Kavehpour, H. P., Giza, C. C., & Prins, M. L. (2023). Traumatic brain injury and the pathways to cerebral tau accumulation. Frontiers in Neurology, 14, Article 1239653. 10.3389/fneur.2023.1239653

28. Hall, C. N., Reynell, C., Gesslein, B., Hamilton, N. B., Mishra, A., Sutherland, B. A., O’Farrell, F. M., Buchan, A. M., Lauritzen, M., & Attwell, D. (2014). Capillary pericytes regulate cerebral blood flow in health and disease. Nature, 508(7494), 55–60. 10.1038/nature13165

29. Harding, A. J., Kril, J. J., & Halliday, G. M. (2000). Practical measures to simplify the Braak tangle staging method for routine pathological screening. Acta Neuropathologica, 99(2), 199–208. 10.1007/PL00007425

30. Hicks, A. J., Sinclair, B., Shultz, S. R., Pham, W., Silbert, L. C., Schwartz, D. L., Rowe, C. C., Ponsford, J. L., Law, M., & Spitz, G. (2023). Associations of Enlarged Perivascular Spaces With Brain Lesions, Brain Age, and Clinical Outcomes in Chronic Traumatic Brain Injury. Neurology, 101(1), e63–e73. 10.1212/WNL.0000000000207370

31. Hussong, S. A., Banh, A. Q., Van Skike, C. E., Dorigatti, A. O., Hernandez, S. F., Hart, M. J., Ferran, B., Makhlouf, H., Gaczynska, M., Osmulski, P. A., McAllen, S. A., Dineley, K. T., Ungvari, Z., Perez, V. I., Kayed, R., & Galvan, V. (2023). Soluble pathogenic tau enters brain vascular endothelial cells and drives cellular senescence and brain microvascular dysfunction in a mouse model of tauopathy. Nature Communications, 14, Article 2367. 10.1038/s41467-023-37840-y

32. Iadecola, C. (2013). The pathobiology of vascular dementia. Neuron, 80(4), 844–866. 10.1016/j.neuron.2013.10.008

33. Iliff, J. J., Wang, M., Liao, Y., Plogg, B. A., Peng, W., Gundersen, G. A., Benveniste, H., Vates, G. E., Deane, R., Goldman, S. A., Nagelhus, E. A., & Nedergaard, M. (2012). A paravascular pathway facilitates CSF flow through the brain parenchyma and the clearance of interstitial solutes, including amyloid β. Science Translational Medicine, 4(147), Article 147ra111. 10.1126/scitranslmed.3003748

34. Iliff, J. J., Chen, M. J., Plog, B. A., Zeppenfeld, D. M., Soltero, M., Yang, L., Singh, I., Deane, R., & Nedergaard, M. (2014). Impairment of glymphatic pathway function promotes tau pathology after traumatic brain injury. The Journal of Neuroscience, 34(49), 16180–16193. 10.1523/JNEUROSCI.3020-14.2014

35. Iverson, G. L., Gardner, A. J., Shultz, S. R., Solomon, G. S., McCrory, P., Zafonte, R., Perry, G., Hazrati, L. -. N., Keene, C. D., & Castellani, R. J. (2019). Chronic traumatic encephalopathy neuropathology might not be inexorably progressive or unique to repetitive neurotrauma. Brain, 142(12), 3672–3693. 10.1093/brain/awz286

36. Jung, L. B., Wiegand, T. L. T., Tuz-Zahra, F., Tripodis, Y., Iliff, J. J., Piantino, J., Arciniega, H., Kim, C. L., Pankatz, L., Bouix, S., Lin, A. P., Alosco, M. L., Daneshvar, D. H., Mez, J., Sepehrband, F., Rathi, Y., Pasternak, O., Coleman, M. J., Adler, C. H., … Koerte, I. K. (2024). Repetitive head impacts and perivascular space volume in former American football players. JAMA Network Open, 7(8), Article e2428687. 10.1001/jamanetworkopen.2024.28687

37. Kaur, R., Singh, V., Kumari, P., Singh, R., Chopra, H., & Emran, T. B. (2022). Novel insights on the role of VCAM-1 and ICAM-1: Potential biomarkers for cardiovascular diseases. Annals of Medicine and Surgery, 84, Article 104802. 10.1016/j.amsu.2022.104802

38. Kim, H. G., Shin, N. Y., Nam, Y., Yun, E., Yoon, U., Lee, H. S., & Ahn, K. J. (2023). MRI-visible Dilated Perivascular Space in the Brain by Age: The Human Connectome Project. Radiology, 306(3), e213254. 10.1148/radiol.213254

39. Kirsch, D., Shah, A., Dixon, E., Kelley, H., Cherry, J. D., Xia, W., Daley, S., Aytan, N., Cormier, K., Kubilus, C., Mathias, R., Alvarez, V. E., Huber, B. R., McKee, A. C., & Stein, T. D. (2023). Vascular injury is associated with repetitive head impacts and tau pathology in chronic traumatic encephalopathy. Journal of Neuropathology & Experimental Neurology, 82(2), 127–139. 10.1093/jnen/nlac122

40. Lau, K., Kotzur, R. & Richter, F. Blood–brain barrier alterations and their impact on Parkinson’s disease pathogenesis and therapy. Transl Neurodegener 13, 37 (2024). 10.1186/s40035-024-00430-z

41. Lin, F., Yang, B., Chen, Y., Zhao, W., Li, B., & Jia, W. (2022). Enlarged perivascular spaces are linked to freezing of gait in Parkinson’s disease. Frontiers in Neurology, 13, Article 985294. 10.3389/fneur.2022.985294

42. Lynch, C. E., Crynen, G., Ferguson, S., Mouzon, B., Paris, D., Ojo, J., Leary, P., Crawford, F., & Bachmeier, C. (2016). Chronic cerebrovascular abnormalities in a mouse model of repetitive mild traumatic brain injury. Brain Injury, 30(12), 1414–1427. 10.1080/02699052.2016.1219060

43. Lynch, K. M., Sepehrband, F., Toga, A. W., & Choupan, J. (2023). Brain perivascular space imaging across the human lifespan. NeuroImage, 271, 120009. 10.1016/j.neuroimage.2023.120009

44. McAllister D, Akers C, Boldt B, Mitchell LA, Tranvinh E, Douglas D, Goubran M, Rosenberg J, Georgiadis M, Karimpoor M, DiGiacomo P, Mouchawar N, Grant G, Camarillo D, Wintermark M and Zeineh MM (2021) Neuroradiologic Evaluation of MRI in High-Contact Sports. Front. Neurol. 12:701948. 10.1089/neu.2023.0505

45. McCann, H., Bahar, A. Y., Burkhardt, K., Gardner, A. J., Halliday, G. M., Iverson, G. L., & Shepherd, C. E. (2022). Prevalence of chronic traumatic encephalopathy in the Sydney Brain Bank. Brain communications, 4(4), fcac189. 10.1093/braincomms/fcac189

46. McKee, A. C., Mez, J., Abdolmohammadi, B., Butler, M., Huber, B. R., Uretsky, M., Babcock, K., Cherry, J. D., Alvarez, V. E., Martin, B., Tripodis, Y., Palmisano, J. N., Cormier, K. A., Kubilus, C. A., Nicks, R., Kirsch, D., Mahar, I., McHale, L., Nowinski, C., … Alosco, M. L. (2023). Neuropathologic and clinical findings in young contact sport athletes exposed to repetitive head impacts. JAMA Neurology, 80(10), 1037–1050. 10.1001/jamaneurol.2023.2907

47. McKee, A. C., Stein, T. D., Huber, B. R., Crary, J. F., Bieniek, K., Dickson, D., Alvarez, V. E., Cherry, J. D., Farrell, K., Butler, M., Uretsky, M., Abdolmohammadi, B., Alosco, M. L., Tripodis, Y., Mez, J., & Daneshvar, D. H. (2023). Chronic traumatic encephalopathy (CTE): Criteria for neuropathological diagnosis and relationship to repetitive head impacts. Acta Neuropathologica, 145(4), 371–394. 10.1007/s00401-023-02540-w

48. Mestre, H., Tithof, J., Du, T., Song, W., Peng, W., Sweeney, A. M., Olveda, G., Thomas, J. H., Nedergaard, M., & Kelley, D. H. (2018). Flow of cerebrospinal fluid is driven by arterial pulsations and is reduced in hypertension. Nature Communications, 9, Article 4878. 10.1038/s41467-018-07318-3

49. Murray, H. C., Osterman, C., Bell, P., Vinnell, L., & Curtis, M. A. (2022). Neuropathology in chronic traumatic encephalopathy: A systematic review of comparative post-mortem histology literature. Acta Neuropathologica Communications, 10, Article 108. 10.1186/s40478-022-01413-9

50. Ojo, J. O., Mouzon, B., Algamal, M., Leary, P., Lynch, C., Abdullah, L., Evans, J., Mullan, M., Bachmeier, C., Stewart, W., & Crawford, F. (2016). Chronic repetitive mild traumatic brain injury results in reduced cerebral blood flow, axonal injury, gliosis, and increased T-tau and tau oligomers. Journal of Neuropathology & Experimental Neurology, 75(7), 636–655. 10.1093/jnen/nlw035

51. Osterman, C., Hamlin, D., Suter, C. M., Affleck, A. J., Gloss, B. S., Turner, C. P., Faull, R. L. M., Stein, T. D., McKee, A. C., Buckland, M. E., Curtis, M. A., & Murray, H. C. (2025). Perivascular glial reactivity is a feature of phosphorylated tau lesions in chronic traumatic encephalopathy. Acta Neuropathologica, 149(1), Article 16. 10.1007/s00401-025-02854-x

52. Plá, V., Bitsika, S., Giannetto, M.J. et al. Structural characterization of SLYM—a 4th meningeal membrane. Fluids Barriers CNS 20, 93 (2023). 10.1186/s12987-023-00500-w

53. Querzola, G., Lovati, C., Laganà, M. M., Pirastru, A., Baglio, F., & Pantoni, L. (2022). Incipient chronic traumatic encephalopathy in active American football players: Neuropsychological assessment and brain perfusion measures. Neurological Sciences, 43(9), 5383–5390. 10.1007/s10072-022-06212-7

54. Ramirez, J., Berberian, S. A., Breen, D. P., Gao, F., Ozzoude, M., Adamo, S., Scott, C. J. M., Berezuk, C., Yhap, V., Mestre, T. A., Marras, C., Tartaglia, M. C., Grimes, D., Jog, M., Kwan, D., Tan, B., Binns, M. A., Arnott, S. R., Bartha, R., … Lang, A. E. (2022). Small and large magnetic resonance imaging-visible perivascular spaces in the basal ganglia of Parkinson’s disease patients. Movement Disorders, 37(6), 1304–1309. 10.1002/mds.29010

55. Rosen, G., Kirsch, D., Horowitz, S., Cherry, J. D., Nicks, R., Kelley, H., Uretsky, M., Dell’Aquila, K., Mathias, R., Cormier, K. A., Kubilus, C. A., Mez, J., Tripodis, Y., Stein, T. D., Alvarez, V. E., Alosco, M. L., McKee, A. C., & Huber, B. R. (2023). Three-dimensional evaluation of cerebrovascular density and branching in chronic traumatic encephalopathy. Acta Neuropathologica Communications, 11(1), Article 123. 10.1186/s40478-023-01612-y

56. Sengillo, J. D., Winkler, E. A., Walker, C. T., Sullivan, J. S., Johnson, M., & Zlokovic, B. V. (2013). Deficiency in mural vascular cells coincides with blood–brain barrier disruption in Alzheimer’s disease. Brain Pathology, 23(3), 303–310. 10.1111/bpa.12004

57. Shepherd, C. E., McCann, H., McLean, C. A., Iverson, G. L., & Gardner, A. J. (2024). Chronic traumatic encephalopathy neuropathologic change in former Australian rugby players. Neuropathology and Applied Neurobiology, 50(2), Article e12972. 10.1111/nan.12972

58. Shepherd, C. E., McGeachie, A. B., Affleck, A. J., & Halliday, G. M. (2019). Condensing the Aβ protocol to reduce the effort and cost of NIA-AA guidelines for neuropathologic assessment of Alzheimer disease. Journal of Neuropathology & Experimental Neurology, 78(10), 975–977. 10.1093/jnen/nlz069

59. Standring, O. J., Friedberg, J., Tripodis, Y., Chua, A. S., Cherry, J. D., Alvarez, V. E., Huber, B. R., Xia, W., Mez, J., Alosco, M. L., Nicks, R., Mahar, I., Pothast, M. J., Gardner, H. M., Meng, G., Palmisano, J. N., Martin, B. M., Dwyer, B., Kowall, N. W., … Stein, T. D. (2019). Contact sport participation and chronic traumatic encephalopathy are associated with altered severity and distribution of cerebral amyloid angiopathy. Acta Neuropathologica, 138(3), 401–413. 10.1007/s00401-019-02031-x

60. Stein, T. D., Alvarez, V. E., & McKee, A. C. (2014). Chronic traumatic encephalopathy: A spectrum of neuropathological changes following repetitive brain trauma in athletes and military personnel. Alzheimer’s Research & Therapy, 6, Article 4. 10.1186/alzrt234

61. Steinman, J., Sun, H. S., & Feng, Z. P. (2021). Microvascular alterations in Alzheimer’s disease. Frontiers in Cellular Neuroscience, 14, Article 618986. 10.3389/fncel.2020.618986

62. Suter, C. M., Cropley, J. E., Affleck, A. J., Lee, M., Hammond, K., Gloss, B., & Buckland, M. E. (2025). Spatially resolved transcriptomics reveals a unique disease signature and potential biomarkers for chronic traumatic encephalopathy. Journal of Neuropathology & Experimental Neurology, 84(11), 967–977. 10.1093/jnen/nlaf078

63. Taoka, T., Ito, R., Nakamichi, R., Nakane, T., Kawai, H., & Naganawa, S. (2024). Diffusion tensor image analysis along the perivascular space (DTI-ALPS): Revisiting the meaning and significance of the method. Magnetic Resonance in Medical Sciences, 23(3), 268–290. 10.2463/mrms.rev.2023-0175

64. Tu, Y., Zhuo, W., Peng, J., Huang, R., Li, B., Liu, Y., Zhang, C., Zeng, X., & Huang, L. (2022). The correlation between enlarged perivascular spaces and cognitive impairment in Parkinson’s disease and vascular parkinsonism. BMC Neurology, 22, Article 282. 10.1186/s12883-022-02819-7

65. Turk, K. W., Geada, A., Alvarez, V. E., Xia, W., Cherry, J. D., Nicks, R., Meng, G., Daley, S., Tripodis, Y., Huber, B. R., Budson, A. E., Dwyer, B., Kowall, N. W., Cantu, R. C., Goldstein, L. E., Katz, D. I., Stern, R. A., Alosco, M. L., Mez, J., … Stein, T. D. (2022). A comparison between tau and amyloid-β cerebrospinal fluid biomarkers in chronic traumatic encephalopathy and Alzheimer disease. Alzheimer’s Research & Therapy, 14, Article 28. 10.1186/s13195-022-00976-y

66. Wardlaw, J. M., Benveniste, H., Nedergaard, M., Zlokovic, B. V., Mestre, H., Lee, H., Doubal, F. N., Brown, R., Ramirez, J., MacIntosh, B. J., Tannenbaum, A., Ballerini, L., Rungta, R. L., Boido, D., Sweeney, M., Montagne, A., Charpak, S., Joutel, A., Smith, K. J., & Black, S. E. (2020). Perivascular spaces in the brain: Anatomy, physiology and pathology. Nature Reviews Neurology, 16(3), 137–153. 10.1038/s41582-020-0312-z

67. Winkler, E. A., Bell, R. D., & Zlokovic, B. V. (2011). Central nervous system pericytes in health and disease. Nature neuroscience, 14(11), 1398–1405. 10.1038/nn.2946

68. Yamamoto EA, Koike S, Luther M, et al. Perivascular Space Burden and Cerebrospinal Fluid Biomarkers in US Veterans With Blast-Related Mild Traumatic Brain Injury. Journal of Neurotrauma. 2024;41(13-14):1565–1577. 10.3389/fneur.2021.701948

69. Yang, P., Pavlovic, D., Waldvogel, H., Dragunow, M., Synek, B., Turner, C., Faull, R., & Guan, J. (2015). String vessel formation is increased in the brain of Parkinson disease. Journal of Parkinson’s Disease, 5(4), 821–836. 10.3233/JPD-140454

70. Zeppenfeld, D. M., Simon, M., Haswell, J. D., D’Abreo, D., Murchison, C., Quinn, J. F., Grafe, M. R., Woltjer, R. L., Kaye, J., & Iliff, J. J. (2017). Association of perivascular localization of aquaporin-4 with cognition and Alzheimer disease in aging brains. JAMA Neurology, 74(1), 91–99. 10.1001/jamaneurol.2016.4370

71. Zhuo J, Raghavan P, Li J, Roys S, Njonkou Tchoquessi RL, Chen H, Wickwire EM, Parikh GY, Schwartzbauer GT, Grattan LM, Wang Z, Gullapalli RP and Badjatia N (2024) Longitudinal assessment of glymphatic changes following mild traumatic brain injury: Insights from perivascular space burden and DTI-ALPS imaging. Front. Neurol. 15:1443496. doi: 10.3389/fneur.2024.1443496

