## Supplementary material for "Neuropathological assessment of cerebrovascular integrity in Chronic Traumatic Encephalopathy"

**Table S1: Pathologies in CTE-NC cases**

| Case number | Co-pathologies |
| --- | --- |
| 1 | CTE-NC, high AD-NC, LBD Braak stage V, LATE-NC stage 2 |
| 2 | CTE-NC, LBD Braak stage VI |
| 3 | CTE-NC, hippocampal sclerosis |
| 4 | CTE-NC, hippocampal sclerosis |
| 5 | CTE-NC, intermediate AD-NC, LATE-NC stage 2 |
| 6 | CTE-NC, high AD-NC, LATE-NC stage 2 |
| 7 | CTE-NC, LATE-NC stage 3 |
| 8 | CTE-NC, LBD Braak stage V, hippocampal sclerosis, LATE-NC stage 3 |
| 9 | CTE-NC, intermediate AD-NC, LBD Braak stage IV, LATE-NC stage 3 |
| 10 | CTE-NC, high AD-NC, amygdala predominant LBD, LATE-NC stage 3 |
| 11 | CTE-NC, amygdala predominant LBD, LATE-NC stage 2 |

CTE-NC = chronic traumatic encephalopathy neuropathological change, AD-NC = Alzheimer’s disease neuropathological change, LBD = Lewy body disease, LATE = Limbic age related TDP-43 encephalopathy, FTLD-TDP = frontotemporal lobar degeneration with TDP-43 immunoreactivity pathology.

**Table S2: Group means and significance**

|  | Control | | | CTE | | | |
| --- | --- | --- | --- | --- | --- | --- | --- |
|  | **Mean** | **SDev** | **n** | **Mean** | **SDev** | **n** | ***p-value*** |
| Arteriolar width (GM) (µm) | 12.8 | 1.41 | 8 | 12.6 | 1.41 | 8 | 0.795 |
| Venular width (GM) (µm) | 10.0 | 1.70 | 8 | 11.2 | 1.70 | 8 | 0.353 |
| Total PVS width (GM) (µm) | 22.7 | 3.11 | 8 | 23.8 | 3.11 | 8 | 0.543 |
| Arteriolar width (WM) (µm) | 15.4 | 3.39 | 8 | 13.2 | 3.39 | 8 | 0.252 |
| Venular width (WM) (µm) | 9.0 | 1.98 | 8 | 9.3 | 1.98 | 8 | 0.805 |
| Total PVS width (WM) (µm) | 24.4 | 5.09 | 8 | 22.5 | 5.09 | 8 | 0.468 |
| Arteriolar to venular PVS distance ratio (WM) | 1.7 | 0.28 | 8 | 1.4 | 0.28 | 8 | 0.086 |
| Arteriolar to venular PVS distance ratio (GM) | 1.3 | 0.28 | 8 | 1.2 | 0.28 | 8 | 0.191 |
| PVS to lumen area ratio (SD) | 3.8 | 1.32 | 7 | 4.4 | 1.32 | 7 | 0.454 |
| PVS to lumen area ratio (GC) | 4.0 | 1.98 | 8 | 5.1 | 1.98 | 8 | 0.307 |
| AQP4 ratio | 1.4 | 0.49 | 6 | 1.6 | 0.28 | 8 | 0.538 |
| Total capillary length (µm) | 2334.14 | 384.38 | 8 | 2418.76 | 384.38 | 8 | 0.693 |
| String vessel % | 0.5 | 0.57 | 8 | 1.0 | 0.57 | 8 | 0.128 |
| Pericyte density (pericytes per µm capillary length) | 7.4 | 3.11 | 8 | 6.4 | 3.11 | 8 | 0.589 |

Group comparisons were assessed using univariate and multivariate regression analyses, as appropriate.

WM = white matter, GM = grey matter, PVS = perivascular space, AQP4 = aquaporin-4, SDev = standard deviation.

**Table S3: Antibodies and reagents used in the study**

| **Antibody/Reagent** | **Species** | **Target** | **Dilution for IHC** | **Company (Catalog #)** | **Application** |
| --- | --- | --- | --- | --- | --- |
| Collagen IV | Goat | Basement  membrane | 1:500 | Novus Biologicals  (NBP1-26549) | SV, Pericyte |
| GFAP | Goat | Astrocytes | 1:200 | My Bio Source  (MBS241916) | PVS |
| SMA-Cy3 | Mouse | Smooth muscle | 1:10000 | Sigma-Aldrich  (C6198) | PVS |
| Aquaporin-4 | Rabbit | Aquaporin-4 | 1:100 | Novus Biologicals  (NBP1-87679) | AQP4 |
| Factor VIII/Von  Willebrand Factor  Phospho-tau (AT8) | Rabbit  Mouse | Endothelial cells  Tau pathology | 1: 10000  1:1500 | Dako (A008202)  Invitrogen (MN1020) | PVS  Neurofibrillary tangles |
| Anti-goat Cy2  secondary antibody | Donkey |  | 1:200 | Jackson  ImmunoResearch  (705-225-147) | PVS |
| Anti-goat Alexa 488  secondary antibody | Donkey |  | 1:200 | Invitrogen (A-  10037) | Pericyte |
| Anti-rabbit Alexa 647  secondary antibody | Donkey |  | 1:200 | Invitrogen (A-  31573) | PVS |
| Anti-goat biotinylated  secondary antibody | Horse |  | 1:200 | Vector Laboratories  (BA9500) | SV |
| VECTASTAIN Elite ABCPeroxidase Kit | Universal |  | 1:500 | Vector Laboratories  (PK6100) | SV |
| 3,3’-diaminobenzidine  (DAB) | Universal |  | 1:33 | Vector Laboratories  (SK4105) | SV |
| TrueBlack Lipofuscin  Autofluorescence  Quencher | Universal |  | 1:20 | Biotium (23007) | PVS, Pericyte |
| DAPI Nucleic Acid Stain | Universal |  | 1:1000 | Invitrogen (D1306) | PVS, Pericyte, AQP4 |

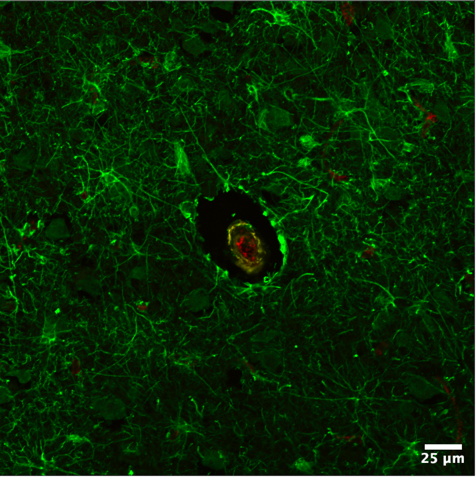

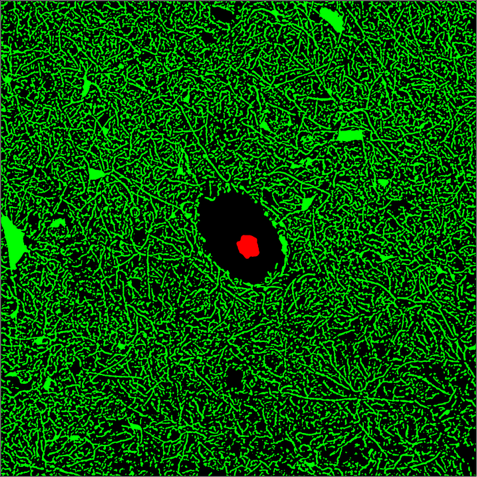

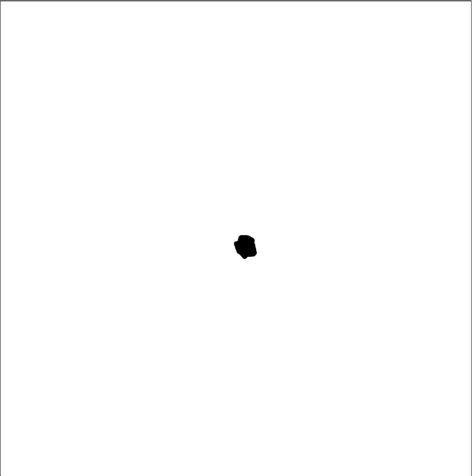

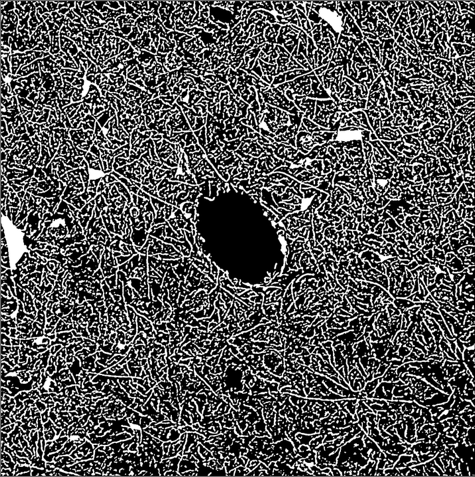

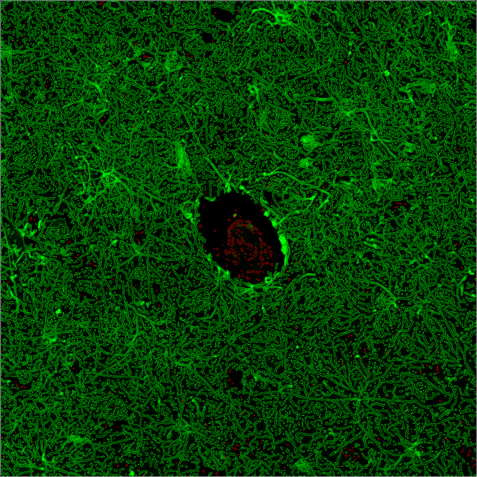

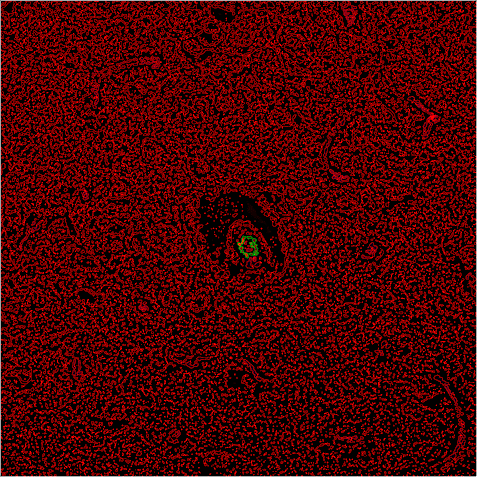

**B**

**A**

**C**

**D**

**E**

**F**

**Supplementary Figure 1:** Micrographs demonstrating the workflow undertaken for PVS quantitation. **(A)** An arteriolar blood vessel from a tissue section from the middle frontal gyrus stained with GFAP (green; astrocytes, white arrow), smooth muscle actin (yellow; smooth muscle, yellow arrow) and factor VIII (red; endothelium, red arrow). **(B)** The endothelial layer (white arrow) and outer perivascular border **(C,** white arrow**)** were identified through fluorescence intensity as the key objects of interest in this analysis. Masks were created to demonstrate the inner **(D)** and outer **(E)** boundaries of the blood vessel. These masks were merged **(F)** and the mean width (μm) between the two was obtained.

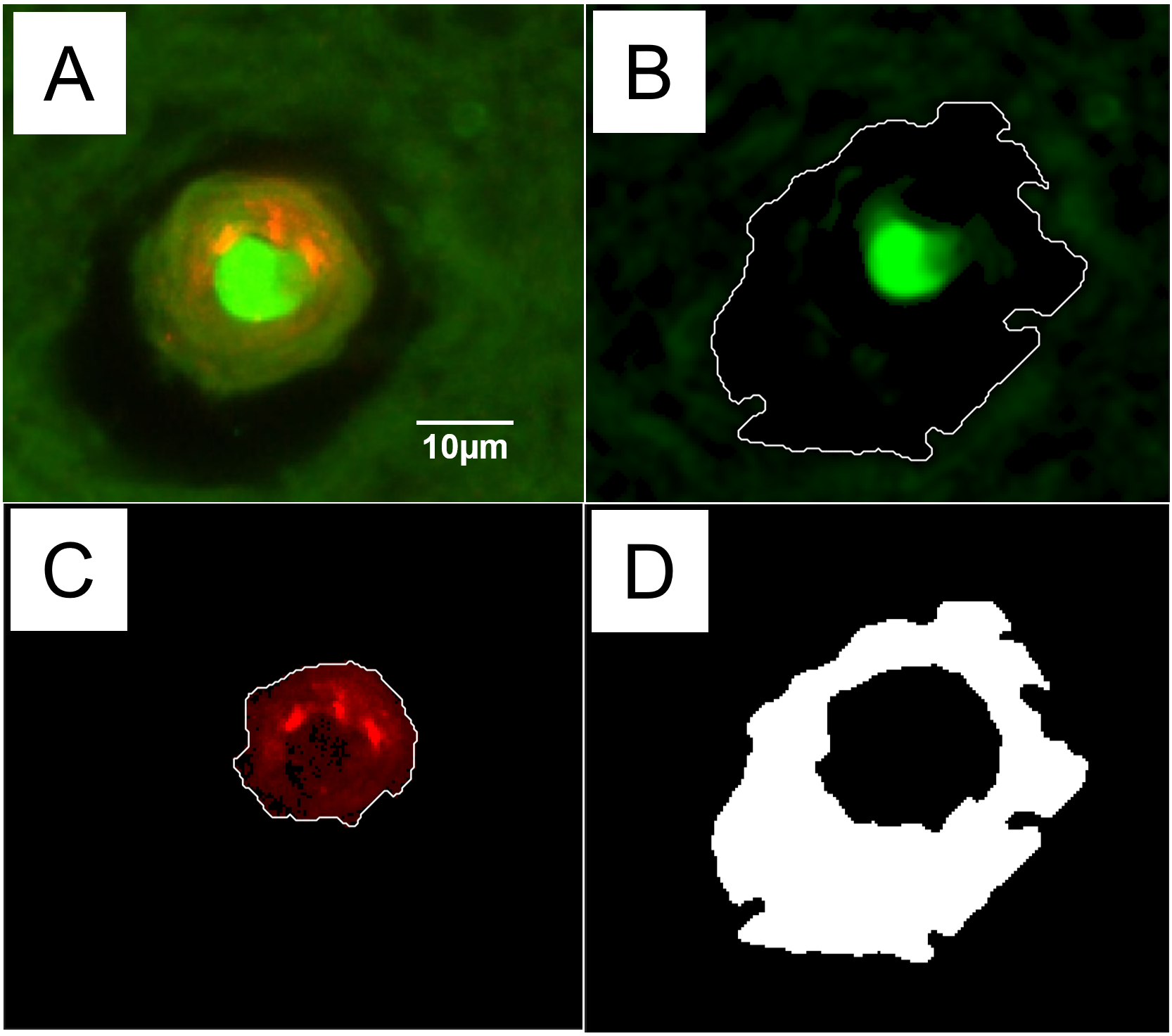

**Supplementary Figure 2. Threshold-assisted manual segmentation for area-based PVS analysis. (A)** Representative vessel-centred ROI showing the green fluorescence channel, containing GFAP immunoreactivity together with prominent tissue autofluorescence, surrounding a factor VIII-positive vessel (red); scale bar = 10 µm. **(B)** Processed green channel used to assist manual delineation of the outer perivascular boundary, shown in white. **(C)** Processed factor VIII channel used to assist manual delineation of the vessel lumen, shown in white. Background correction and adaptive thresholding were used to improve discrimination of the relevant fluorescence boundaries; preprocessing parameters were adjusted modestly between staining batches to account for differences in fluorescence intensity and background. Final contours were manually defined with reference to the underlying fluorescence signal rather than generated automatically from the thresholded images. **(D)** Final binary PVS mask calculated as the area enclosed by the GFAP-defined outer boundary after subtraction of the vessel lumen.

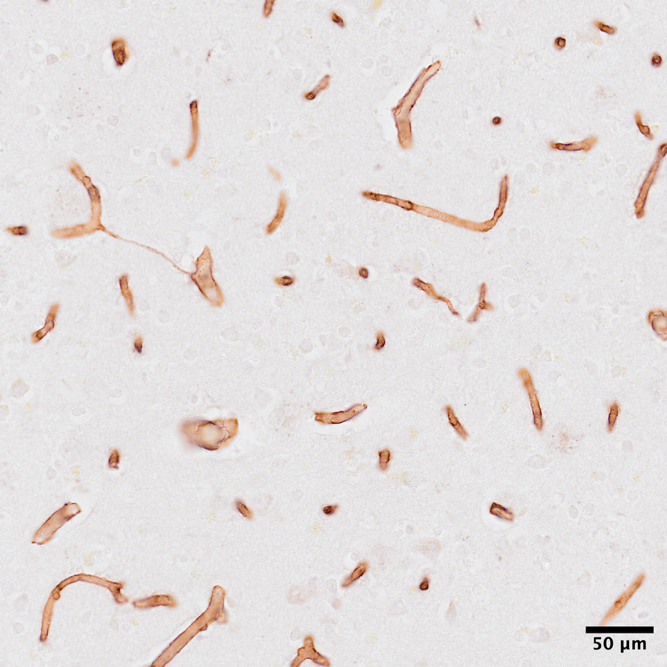

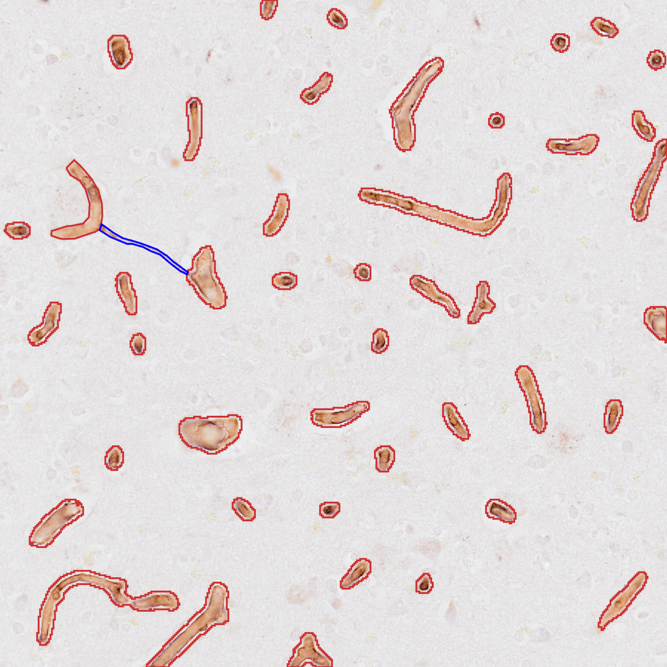
**Supplementary Figure 3:** Representative immunoperoxidase micrographs stained for the detection of the capillary network, including string vessels, in the middle frontal gyrus **(A)**. 5 images were randomly sampled and capillaries were distinguished from background tissue using QuPath **(B**; red outline, black arrow). String vessels were manually segmented and were identified based on their distinctive string-like structure **(B**; blue outline, red arrow). Scale bar for **(A)** is the same for **(B)**.

**B**

**A**

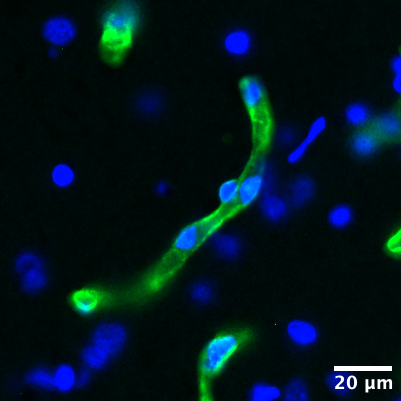

**Supplementary Figure 4:** Representative immunofluorescent micrograph demonstrating pericyte analysis on capillaries in the middle frontal gyrus. Pericyte cell bodies were identified by their morphology, and presence on the outer surface of the capillary wall (white arrow). The pericyte has a DAPI-positive nucleus.

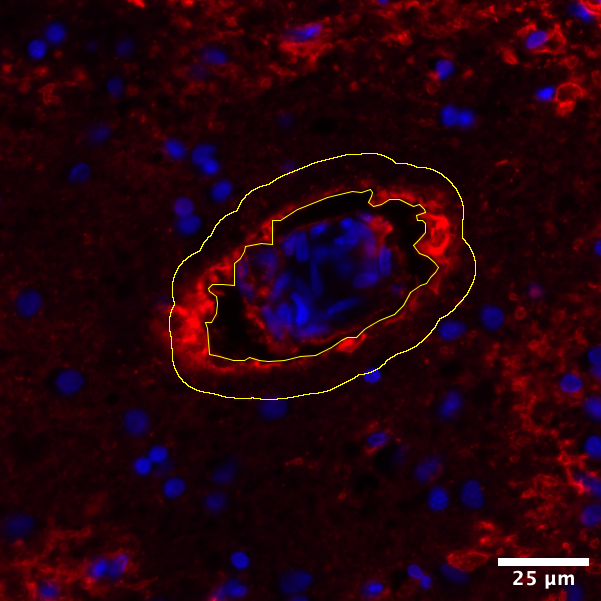

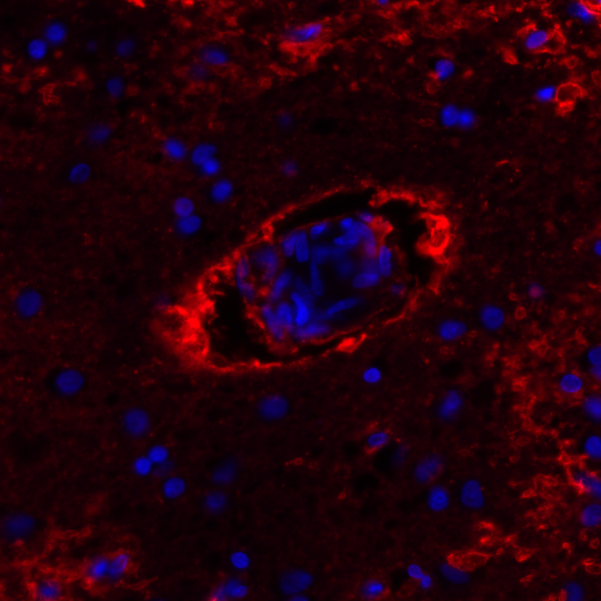

**A**

**B**

**Supplementary Figure 5:** Representative immunofluorescence micrographs demonstrating AQP4 localisation in astrocytic endfeet in the middle frontal gyrus **(A)**. AQP4 localisation around the perivascular border was assessed in 6 regions of interest per case using a 10μm selection band **(B)**. The amount of AQP4 in the astrocytic endfeet surrounding the perivascular border was measured as a ratio of endfeet fluorescence intensity relative to global AQP4 fluorescence intensity (perivascular: global). The scale bar in **(B)** is the same for **(A)**.

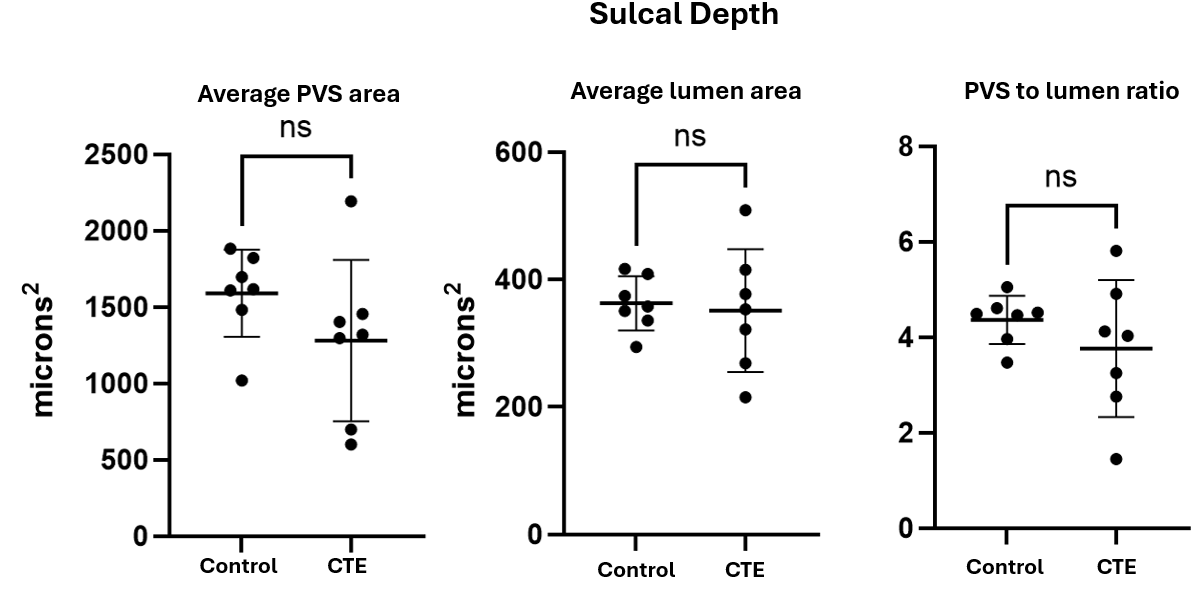

**Supplementary Figure 6.** A comparison of average PVS and lumen area of vessels in the sulcal depth revealed no significant (ns) difference between the control and CTE-NC; Average PVS area (p = 0.097), average lumen area (p = 0.805), PVS to lumen ratio (p = 0.383). In the control samples the mean lumen size recorded was 362.7 ± 42.33 µm², while the mean PVS size was 1590 ± 285.3 µm². In the CTE-NC samples the average lumen size was 351.4 ± 96.5 µm², and the average PVS size was 1281 ± 528.5 µm². The ratio of PVS to lumen areas was 4.4 ± 0.51 in the control case and 3.9 ± 1.2 for CTE-NC. Individual points represent independent cases; horizontal lines and error bars indicate mean ± SD. Mann-Whitney U tests were used, n (CTE-NC) = 7, n (control) = 7.

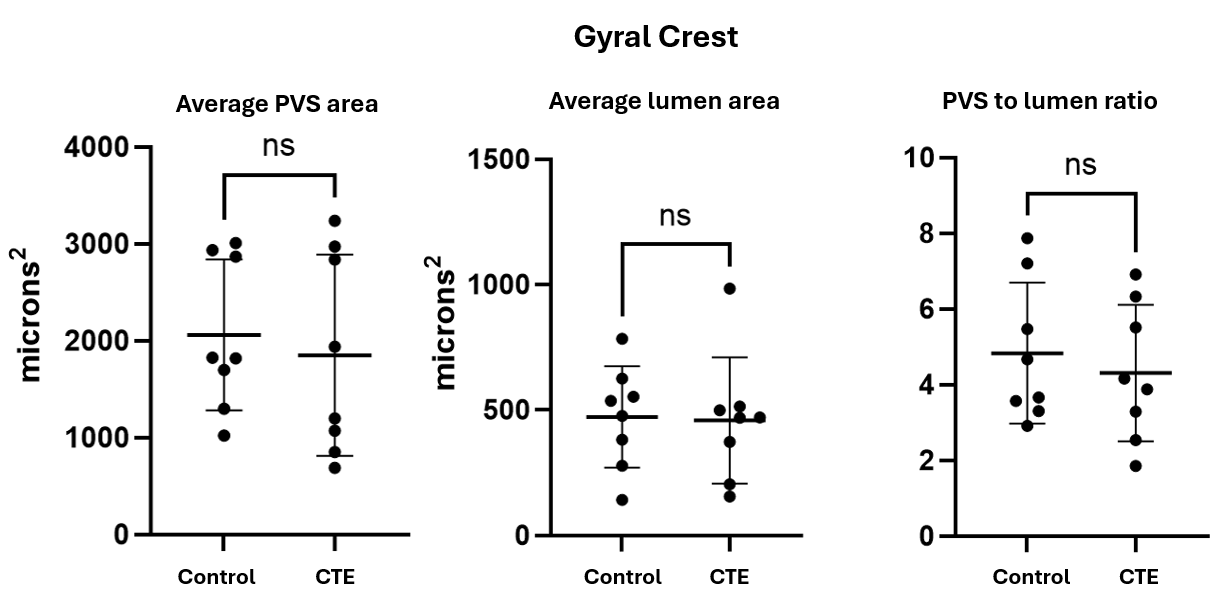

**Supplementary Figure 7.** A comparison of average PVS and lumen area of vessels in the gyral crest revealed no significant (ns) difference between the control and CTE-NC; average PVS area (p = 0.645), average lumen area (p = 0.663), PVS to lumen ratio (p = 0.656). In the control samples the mean lumen size recorded was 472.3 ± 202.2 µm², while the mean PVS size was 2060 ± 777 µm². In the CTE-NC samples the average lumen size was 458.9 ± 252.2 µm², and the average PVS size was 1849 ± 1038 µm². The ratio of PVS to lumen areas was 4.8 ± 1.9 in the control cases and 4.3 ± 1.8 for CTE-NC. Individual points represent independent cases; horizontal lines and error bars indicate mean ± SD. Mann-Whitney U tests were used, n (CTE-NC) = 8, n (control) = 8.

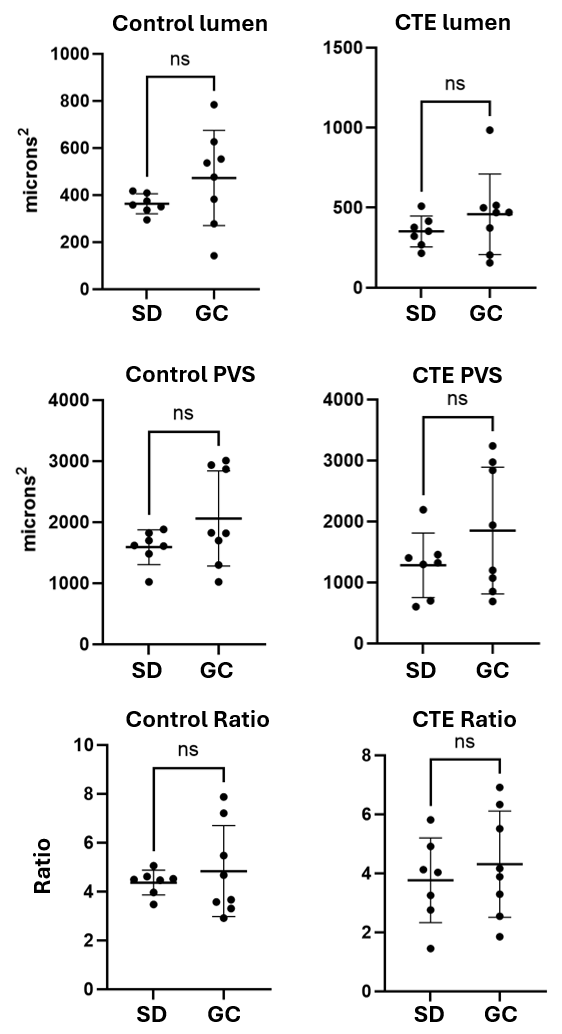

**Supplementary Figure 8:** Comparison between sulcal depth and gyral crests regions for vessel lumen, PVS, and PVS to lumen ratio. There was no significant (ns) difference in any of these measures between regions. Control lumen (p = 0.175), CTE-NC lumen area (p = 0.31), Control PVS (p = 0.156), CTE-NC PVS (p=0.215), Control ratio (p = 0.534), CTE-NC ratio (p=0.531). Individual points represent independent cases; horizontal lines and error bars indicate mean ± SD. Mann-Whitney U tests were used, n (CTE-NC) = 7, n (control) = 7.
